# Functional gradients in the human lateral prefrontal cortex revealed by a comprehensive coordinate-based meta-analysis

**DOI:** 10.1101/2022.01.21.477198

**Authors:** Majd Abdallah, Gaston Zanitti, Valentin Iovene, Demian Wassermann

**Affiliations:** Team Parietal, Inria, CEA, Université Paris-Saclay, Palaiseau, 91120, France; NeuroSpin, CEA, Université Paris-Saclay, Gif-sur-Yvette, 91190, France

**Keywords:** Lateral prefrontal cortex, rostrocaudal gradient, meta-analysis, probabilistic logic programming, neuroinformatics

## Abstract

The human lateral prefrontal cortex (LPFC) enables flexible goal-directed behavior. Yet, its organizing principles remain actively debated despite decades of research. Meta-analysis efforts to map the LPFC have either been restricted in scope or suffered from limited expressivity in meta-analysis tools. The latter short-coming hinders the complexity of questions that can be expressed in a meta-analysis and hence limits the specificity of structure-function associations. Here, we adopt NeuroLang, a novel approach to meta-analysis based on first-order probabilistic logic programming, to infer the organizing principles of the LPFC with greater specificity from 14,371 neuroimaging publications. Our results reveal a rostrocaudal and a dorsoventral gradient, respectively explaining the most and second-most variance in whole-brain meta-analytic connectivity in the LPFC. Moreover, we find a cross-study agreement on a spectrum of increasing abstraction from caudal to rostral LPFC both in specific network connectivity and structure-function associations that supports a domain-general role for the mid-LPFC. Furthermore, meta-analyzing inter-hemispheric asymmetries along the rostrocaudal gradient reveals specific associations with topics of language, memory, response inhibition, and error processing. Overall, we provide a comprehensive mapping of the organizing principles of task-dependent activity in the LPFC, grounding future hypothesis generation on a quantitative overview of past findings.

## 1 Introduction

The human lateral prefrontal cortex (LPFC) supports a wide variety of cognitive processes that are considered hallmark features of the human brain [1, 2]. Understanding the functional organization of the LPFC is thus important to the study of adaptive human behavior. Yet, the overarching organizing principle of the LPFC is still actively debated, with a variety of proposals on whether it is unitary, hierarchical, or houses a set of separable networks subserving distinct functions [3, 4, 5, 6, 7, 8]. There have been a few large-scale attempts to map the entire LPFC, but these mappings often lack specificity, partly due to the limited breadth of queries that common meta-analysis methods can express. In this study, we adopt a novel approach to meta-analysis based on symbolic artificial intelligence to infer the organizing principles of the LPFC from thousands of neuroimaging studies with greater expressivity and specificity.

The versatility of the LPFC suggests that it is far from unitary [5, 6, 9, 10, 11]. An influential class of hypotheses emerging from the domain of abstraction and hierarchical control proposes a rostrocaudal gradient in the LPFC, wherein caudal regions respond to immediate sensory stimuli, middle regions select actions based upon a prevailing context, and rostral regions integrate concrete representations into more abstract rules and goals to enable temporal control of behavior [1, 2, 3, 6, 11, 12, 13, 14, 15, 16]. A second class of hypotheses holds that a dorsoventral gradient segregating regions involved in distinct stimulus domains, such as spatial vs. non-spatial, also governs the distribution of functions in the LPFC [2, 17, 18]. Further results reveal that the ventral, dorsal, and middle LPFC are each organized along their rostrocaudal axes according to the level of abstraction in task representations [2, 19, 20].

Contemporary evidence from systems neuroscience proposes that the LPFC is spanned by distinct functional networks, such as the attention, default mode, and most importantly the salience (SN) and frontoparietal control (FPCN) networks [10, 21, 22, 23, 24]. These networks are globally situated upon a brain-wide intrinsic connectivity gradient, wherein the transmodal regions of the default mode network are maximally distant from sensorimotor unimodal regions [25, 26], with multimodal regions of the SN and FPCN occupying intermediate zones. One proposal holds that this spatial principle ascribes the LPFC with a role in integrating both concrete and abstract representations, suggesting an external/present-oriented to internal/future-oriented gradient extending outwardly from the motor cortex towards the anterior of the brain [8]. However, recent studies that rely on causal evidence argue against a linear unidimensional gradient in the LPFC, and rather support the hypothesis of separable networks dynamically interacting within global and local hierarchies to support adaptive human behavior [6, 7, 9, 10, 27, 28] (also see [29] for a comprehensive review). Within this systems-based framework, the middle LPFC not the rostral LPFC is believed to act as a focal point, integrating concrete and abstract representations from disparate networks, with increasingly rostral and caudal LPFC regions acting in a domain-specific manner [8, 28, 29].

Further, inter-hemispheric functional asymmetries in the LPFC are widely reported, most notably for language [22, 30] and inhibitory control processes [31, 32]. Functional asymmetries between hemispheres are believed to arise from dynamic patterns of inter- and intra-hemispheric connectivity that represent organizing principles of functional specializations whose putative function is to promote efficient control of behavior [33, 34]. Thus, mapping the LPFC should take into account differences across hemispheres, especially in the distribution of lateralized topic associations. While there is a preponderance of research on the organization of the left LPFC in the fields of hierarchical control and language [8], a comprehensive comparison is yet to draw firm conclusions regarding the specifc functional associations of both LPFC hemispheres.

The multitude of proposals on the LPFC may arise from the diversity of protocols and researcher’s degrees-of-freedom in individual studies whose idiosyncrasies (task type, timing, magnitude of stimuli/responses, data analysis methods, and publication bias) can limit generalizability [35, 36]. And besides concerns of small sample sizes [37], each individual study probes a narrow scope of the broad range of functions that putatively engage the LPFC, which poses the risk of interpreting the results based on a small set of task contrasts. Therefore, it remains unclear to what extent the functional boundaries derived from each individual study correspond to the global organization of activity observed in the LPFC during a wider variety of behaviors. Ultimately, a quantitative meta-analysis is needed to make inferences on the global organization of the LPFC. Unlike individual studies, meta-analysis offers an overarching perspective on task-dependent activity and maps a wide range of mental functions onto the LPFC by synthesizing thousands of published findings into a single statistical framework [38, 39, 40].

The few existing meta-analyses on the LPFC, although informative, have been limited in scope, assumptions, and most importantly, tools. These limitations preclude a reliable distinction of closely related LPFC regions in terms of their relative specificity to networks and mental functions. On one hand, due to the difficulty of manual compilation of activation peaks from the literature, most meta-analyses on the LPFC have been restricted to particular regions [e.g. right inferior frontal gyrus 33] or functions [e.g. working memory 41]. On the other hand, large-scale automated meta-analyses [e.g. 9] have assumed that LPFC regions are clusters of piece-wise constant coactivation, ignoring overlaps between them and not specifying an organizing spatial schema of functional transitions from one region to another. Finally, commonly-used tools, such as Neurosynth [40], are not expressive enough to represent complex hypotheses of specific functional associations in the LPFC. For example, it is arguably difficult and arduous to query a meta-analytic database on the probability that a topic is associated with a study given activation in one region and the simultaneous absence of activation in any spatially-anterior region (similarly posterior, superior or homologous). The expressivity limitation becomes challenging when performing a comprehensive meta-analysis with tens of topics at a time as well as regions that are consistently coactivated by several tasks.

Here, we overcome these challenges using a recently-introduced query language, called NeuroLang [42] to perform a comprehensive coordinate-based meta-analysis on 14,371 articles from the Neurosynth database [40], along with a gradient-mapping technique [26] to identify the organizing principles of activity in the LPFC. NeuroLang puts forth first-order probabilistic logic programming as a structured and more expressive formalism to represent neuroscience hypotheses and solve complex queries on large databases [42]. For instance, we can succinctly express queries of functional specificity in the likes of: “*What is the probability that empathy is present given activation in the rostral LPFC and there does not exist any activation reported in caudal or middle LPFC?*”. NeuroLang also brings the power of probabilistic reasoning to deal with elements of uncertainty in heterogeneous data, such as in peak locations, between-regions coactivation, and the presence of terms, all in a single unifying framework. Most importantly, however, a meta-analysis performed using NeuroLang is highly reproducible, that is, the same queries used by one study can be used by future studies to validate the results as more data becomes available.

By leveraging the expressivity of NeuroLang to perform this meta-analysis, we identify a principal rostrocaudal gradient and a subsidiary dorsoventral gradient that respectively explain the most and second-most variance in meta-analytic connectivity in the LPFC in both hemispheres. Moreover, we find that the principal gradient captures a spectrum of increasing abstraction in patterns of network connectivity and specific topic associations from caudal to rostral LPFC, while supporting a proposed domain-general role for middle LPFC regions. Finally, a gradient-based meta-analysis of inter-hemispheric asymmetries reveals the relative dominance of language and memory in the left LPFC as well as the relative dominance of inhibitory control and error processing in the right LPFC.

## 2 Results

### 2.1 Principal rostrocaudal and secondary dorsoventral gradients explain most of the variance in meta-analytic connectivity in the LPFC

In the first analysis, we infer the extent to which LPFC regions agree in the spatial distribution of meta-analytic connectivity patterns across thousands of studies found in the Neurosynth database. In other words, our goal is to identify the main profiles of variation (i.e. gradients) in whole-brain meta-analytic connectivity within the LPFC [25, 26].

Towards achieving this goal, we need to reduce the high-dimensionality of voxel-level data to increase interpretability of our findings and alleviate computational burdens. To do this, we project voxel-level data onto 1024 functional regions from the Dictionaries of Functional Modes (DiFuMo) probabilistic atlases covering the entire brain [43]. The DiFuMo is a set of continuously-valued brain atlases derived from thousands of subjects across 27 studies, including a total of 2192 task-based and resting-state functional magnetic resonance imaging (fMRI) sessions publicly available on OpenNeuro [44]. Unlike spatially-constrained clusters, a DiFuMo atlas does not ignore the overlap among neuronal populations, allowing voxels to be grouped into multiple functional regions with varying weights. To this end, we write NeuroLang logic program (Program available here) that infers the conditional probability of a brain region to be reported active given activation in a LPFC region, as well as the conditional probability that the brain region is active given no activation in the LPFC region. The program then computes the logarithm with base 10 of the odds ratio (LOR) of these two hypotheses for every LPFC-brain region pairs, creating a *N* × *M* meta-analytic connectivity matrix, where *N* denotes the number of regions in the LPFC and *M* the number of regions in the entire brain. The LOR captures the amount of evidence in favor of specific coactivation between each LPFC region and brain regions, as compared to the evidence favoring independent activation of each.

After constructing the coactivation matrix, we apply an unsupervised non-linear dimensionality reduction method known as diffusion embedding [26, 45] to the resultant meta-analytic connectivity matrix in each hemisphere, separately, using the BrainSpace toolbox [46]. The resultant low-dimensional embedding identifies the position of each LPFC region along unidimensional axes, known as gradients, each representing a direction of variation in meta-analytic connectivity. The axis that accounts for the greatest amount of variance in meta-analytic connectivity is called the principal gradient, which will be the focus of the next analyses. Technical details on formulating the NeuroLang logic program that infers the meta-analytic connectivity matrix along with details on diffusion embedding are found in the Materials and Methods section.

The results of this analysis are shown in Figure 1 and Figure 2. Figure 1A depicts the principal gradient of coactivation which explains the greatest percentage of variance in the meta-analytic connectivity in both LPFC hemispheres (see Figure 1C and Figure 1D). This gradient is anchored at one end by caudal LPFC regions and at the other end by rostral LPFC regions, supporting a dominant rostrocaudal organization in the LPFC across the literature. The spatial layout of the principal gradient is expressed in terms of the posterior-to-anterior and inferior-to-superior positions of regions distributed along successive twenty-percentile gradient segments (i.e. quintile bins; see Figure 2). On the other hand, Figure 1B shows the gradient that explains the second-most percentage of variance in connectivity, extending along the dorsoventral axis of the LPFC. This gradient is anchored at on end by ventral LPFC regions and at the other end by dorsal LPFC regions. Collectively, the topographic profiles of the first two gradients of meta-analytic connectivity support and integrate the views that the LPFC is organized along its rostrocaudal and dorsoventral axes. Although the topography of the proposed gradients has been previously described [e.g. 2, 6, 19], the relative extent to which they explain the distribution of activations in the LPFC across a wide variety of brain states have remained unclear. Thus, we contribute by revealing a dominant rostrocaudal gradient representing the overarching organizing principle of task-dependent activation in the LPFC in the literature.

**Figure 1:**
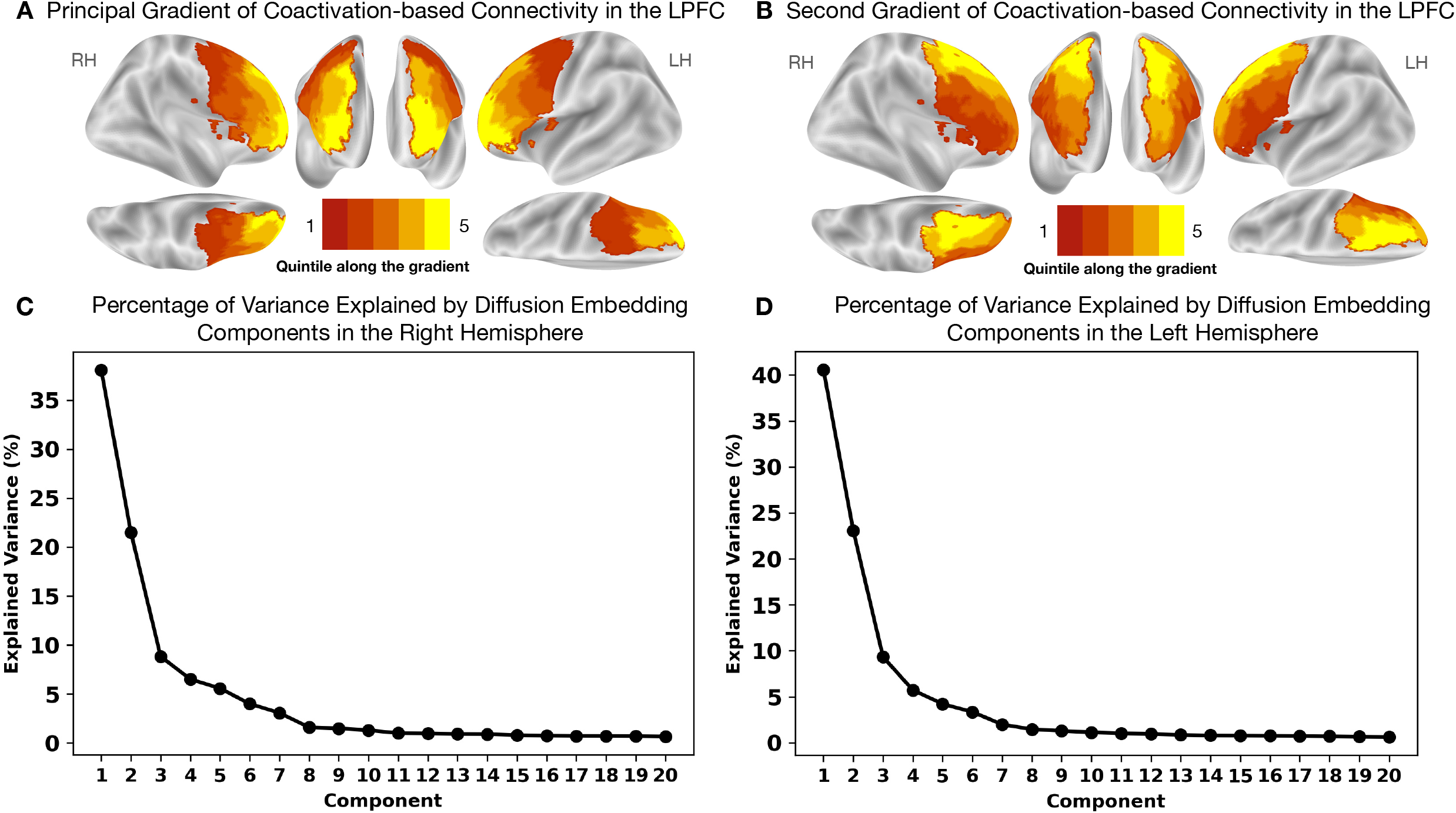
Principal rostrocaudal and secondary dorsoventral gradients explain the greatest amount of variance in meta-analytic connectivity in the LPFC. **(A)** The principal gradient in both hemispheres echoes a widely proposed rostrocaudal organization in the LPFC. This gradient represents the dominant direction of gradual variations in coactivation, and hence function, in the LPFC. **(B)** The gradient that explains the second-most variance in meta-analytic coactivation-based connectivity echoes a dorsoventral organization in the LPFC extending from ventrolateral to dorsolateral PFC regions. **(C)** and **(D)** The amount of variance explained by diffusion embedding components in the right and left LPFC, respectively.

**Figure 2:**
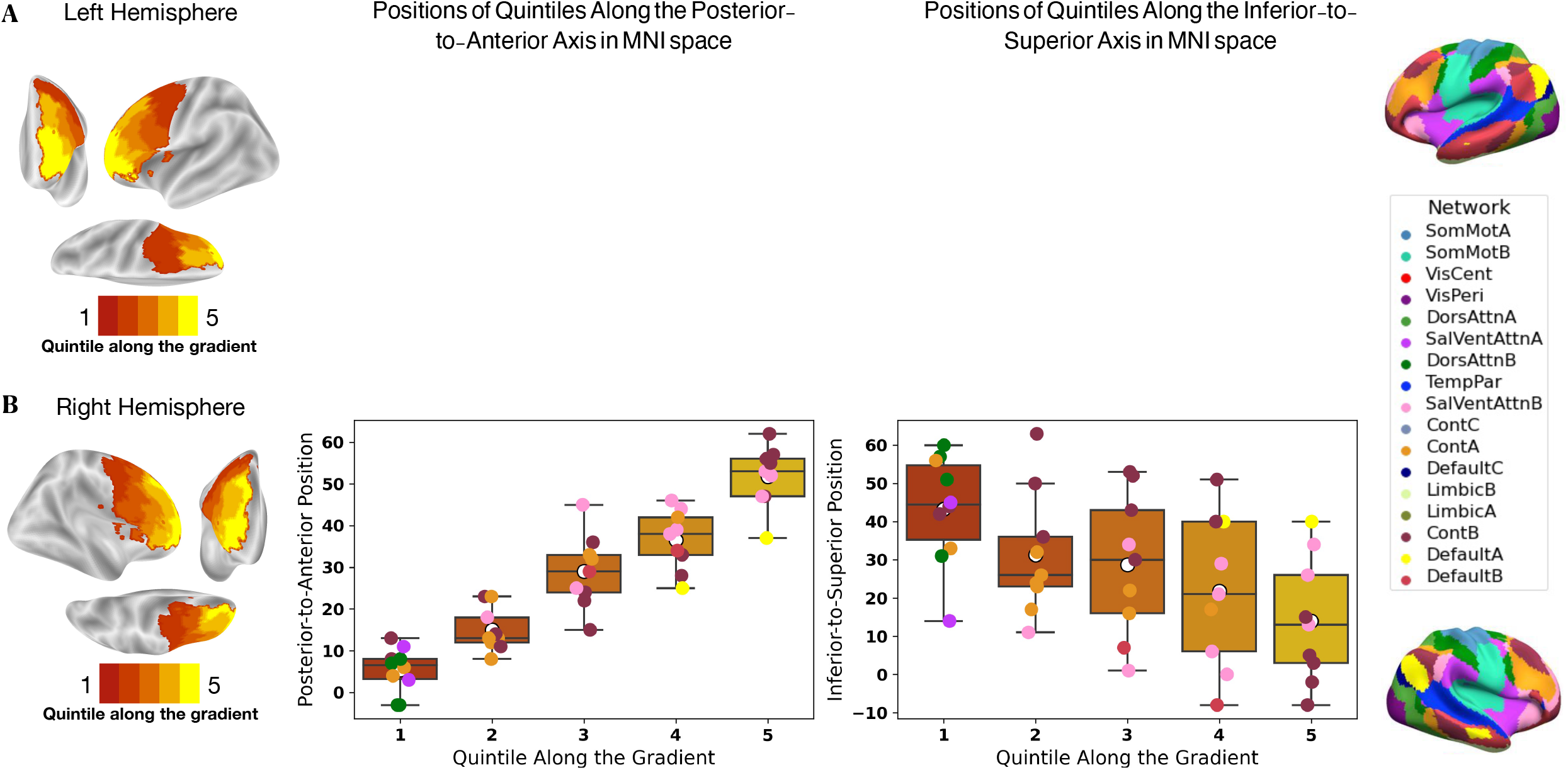
The posterior-to-anterior and inferior-to-superior positions in MNI space of regions grouped into quintile bins along the principal gradient reflect a rostrocaudal organization. **(A)** Positions of regions in the left hemisphere. **(B)** Positions of regions in the right hemisphere. Each colored sphere represents a region in the LPFC, with the color reflecting its network membership within the 17-Networks atlas shown at the left of the figure [24]. **SomMot**: Somatomotor, **VisCent/Peri**: Visual Central/Peripheral, **SalVentAttn**: Salience/Ventral Attention, **DorsAttn**: Dorsal Attention, **TemPar**: Temporo-parietal, **Cont**: Executive Control, **Default**: Default Mode

### 2.2 Varying coactivation patterns along the rostrocaudal gradient capture a spectrum of increasing abstraction in large-scale network connectivity

In the second analysis, we characterize the rostrocaudal gradient in both LPFC hemispheres in terms of varying coactivation patterns of successive twenty-percentile gradient segments (i.e. quintile bins) and their overlap with canonical large-scale brain networks (Figure 3). For this purpose, we write a NeuroLang logic program (Program available here) that first performs a multilevel kernel density analysis [47] using a uniform kernel of 10 mm radius at the study level, and then projects the resulting binary activation maps (1 map per study) onto 1024 functional regions. This yields a mapping between brain regions and each study in the database, wherein each region has a probability of being reported by a study, which depends on the location of the reported voxels (further details are provided in Materials and Methods). However, this is not the case for quintile bins along the principal gradient, which are labels and not continuously valued regions. Therefore, we set the program to consider activation reported in a quintile bin if at least one peak is reported within it or within its near vicinity (< 3*mm*). Consequently, for each quintile bin along the principal gradient, the program infers the logarithmic ratio of odds for a brain region to be reported active given activation in a quintile bin to the odds of the region activation given no bin activation. This ratio represents the amount of evidence for a specific coactivation between a brain region and a given quintile bin along the principal gradient of the LPFC.

**Figure 3:**
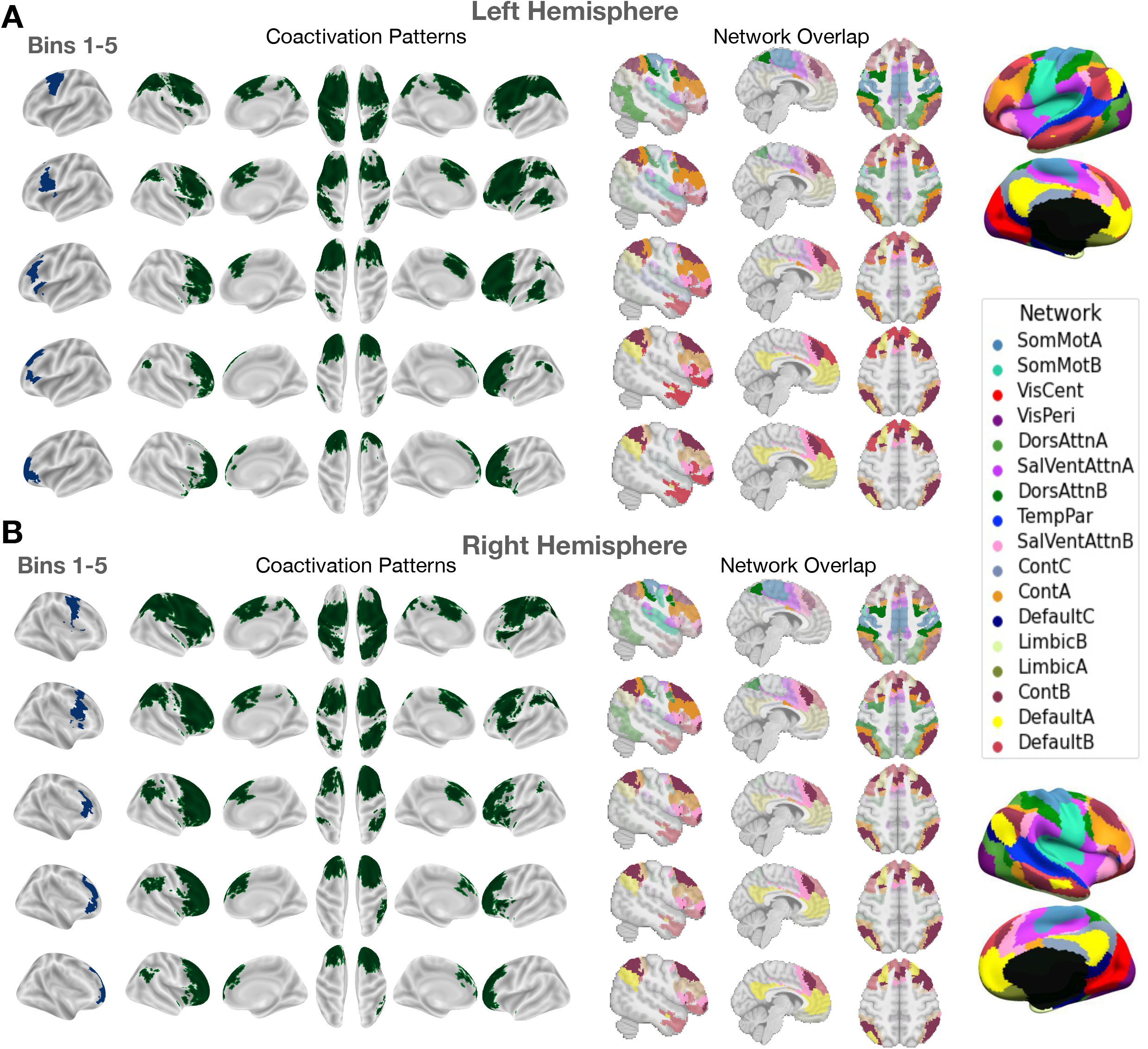
The meta-analytic coactivation patterns of quintile bins along the principal gradient in the LPFC capture a spatial layout of increasing abstraction in canonical network connectivity. **(A)** Coactivation patterns along the principal gradient in the left LPFC. **(B)** Coactivation patterns along the principal gradient in the right LPFC. Each coactivation map shows the regions that have a least three times the odds of being reported active given activation reported in a bin relative to being reported active when no activation is reported in the bin. Note that cerebellar and sub-cortical regions although included in the analysis are not shown in the figures. The **Network Overlap** panel shows the brain networks from the 17-Networks atlas [24] that overlap with the coactivation pattern of each quintile bin. The transparency of the color reflects the relative proportion of volume in the coactivation pattern the overlaps with each brain network. **SomMot**: Somatomotor, **VisCent/Peri**: Visual Central/Peripheral, **SalVentAttn**: Salience/Ventral Attention, **DorsAttn**: Dorsal Attention, **TemPar**: Temporo-parietal, **Cont**: Control, **Default**: Default Mode

The results of this analysis are shown in Figure 3. A cortical coactivation map is constructed by recovering regions that exhibit at least threefold the odds (or *LOR* > 0.5) of being reported active give activation is reported in a bin compared to being active given otherwise. The “Network Overlap” panel in Figure 3A and Figure 3B depicts the large-scale brain networks defined by the 17-Network parcellation [24] that overlap with each bin’s coactivation pattern. The relative proportion of overlap between the coactivation pattern and each network is reflected by the level of color transparency in the brain plot. Increasingly opaque colors indicate that more volume of the coactivation pattern overlaps with a given network, with the most opaque colors signifying predominant networks.

This analysis reveals a structured ordering of network connectivity profiles along the rostrocaudal LPFC gradient in both hemispheres, from a pattern mostly dominated by networks involved in external processing to a pattern dominated by networks involved in internally-oriented cognition. Almost all bins coactivate with the salience and frontoparietal control networks (i.e. SalVentAttnB, ContA and ContB) to varying degrees, with ContA mostly dominating the coactivation pattern of caudal-to-middle zones of the gradient. Moreover, while middle zones’ coactivation patterns mostly overlap with the salience and control networks, they also overlap with both externally and internally focused networks in a rostrocaudal fashion. That is, bin 2 in both hemispheres coactivates more with the dorsal attention networks, while bin 3 coactivates with the default mode network. This result may support contemporary accounts of domain-general integrative processing in the mid-LPFC regions as opposed to more domain-specificity at the extremities of the rostrocaudal LPFC gradient.

### 2.3 Mapping specific topic associations in the LPFC supports the hypothesis of increasing abstract representations extending along the principal rostrocaudal gradient

In the third analysis, we infer specific functional associations of coactivation patterns among quintile bins along the principal gradient of the LPFC in both hemispheres using what we call “segregation queries”. A segregation query infers the probability “*that a topic is present in a study given coactivation in a set of regions and the simultaneous absence of coactivation in another set of regions*”. This type of queries is arguably difficult to express in other automated meta-analysis tools especially as the number of topics and regions increases, but can be readily represented in NeuroLang. Expressing segregation queries using NeuroLang enables the inference of specific structure-functions associations that are otherwise blurred by the coactivation of regions across tasks. The reason for this blurring is that typical fMRI task contrasts rarely isolate regions underlying distinct but related processes, which likely need to be probed across multiple tasks to ensure the independence of regions [48].

We conduct this segregation-based meta-analysis using 38 topics expertly-chosen from an original set of 100 topics (version-5 of topic modelling) from Neurosynth [49]. These topics cover broad cognitive and behavioral domains mainly studied in cognitive neuroscience. We set the NeuroLang logic program (Program available here) to infer the probability *that a topic is present given activation in bin a (a* ∈ [1, 5]*) and bin b (b* ∈ [1, 5]*) and there exists no activation in any bin outside the range* [*a*, *b*]. For instance, in the event where *a* =1 and *b* = 4, the program queries the database on the probability that a topic is present in a study given coactivation of bins 1 and 4 (and potentially any region in between) and there exists no activation in any bin outside the quintile range [1, 4]. In the event where *a* = *b*, the program queries the database on the presence of a topic given activation constrained in only one quintile bin at a time. We illustrate this visually in Figure 4, for example, the coactivation of bins 1 and 4 (and potentially any bins in between) is represented by the “1 < – > 4” notation on the horizontal axis, and the sole activation of bin 3 is represented by the “3” notation. Concurrently, the program infers the probability of the opposite event by selecting the studies that do not match a criteria imposed by a given segregation query. By computing the LOR of the two opposing hypotheses, we obtain a measure of the evidence in favor of specific associations between each topic and spatially-constrained activation patterns along the principal gradient of the LPFC.

**Figure 4:**
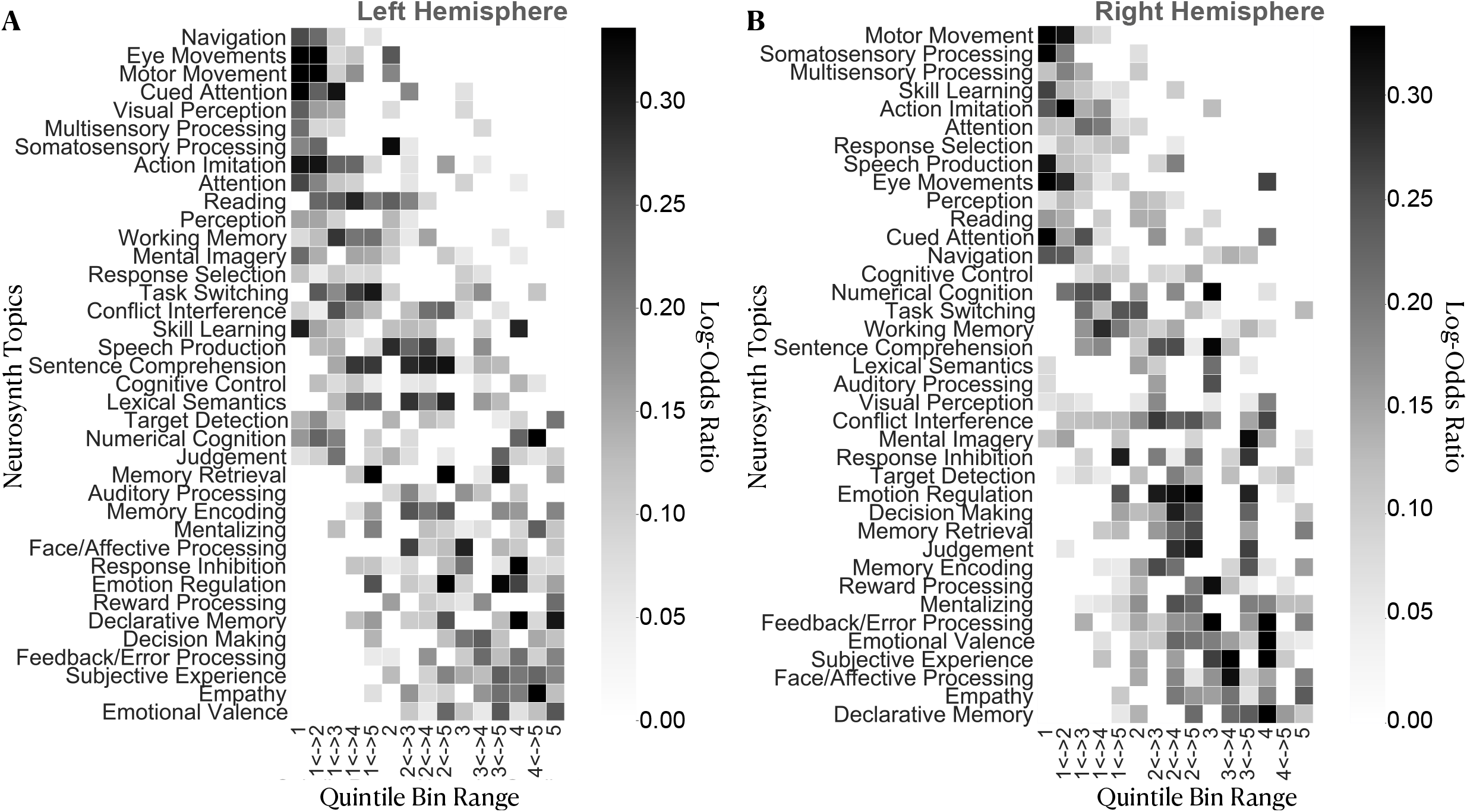
Segregation-based meta-analysis of topic-bin associations using 38 topics reveals a systematic shift in function from caudal to rostral LPFC regions characterized by increasing abstraction. **(A)** and **(B)** show specific topic associations in the left and right hemispheres, respectively. Topics are ordered by the weighted mean of their location along the principal gradient. A two-headed arrow along the horizontal axis signifies a coactivation among quintile bins in a given range and potentially any region within but not outside the range. Note that the exact order of topics varies between hemispheres, but the general profile of topic distribution is comparable.

Results of this analysis are depicted in Figure 4A and Figure 4B for the left and right hemispheres, respectively. Topics are ordered by the weighted mean of their location along the gradient, revealing a systematic shift in topic associations from external processing at the caudal end to more abstract cognitive, affective and memory-related topics at the rostral end of the principal gradient. Between these extremities, we observe domain-general executive functions and topics related to language and semantic processing. This pattern of topic-bin associations suggests that as activation patterns extend away from caudal LPFC (bins 1 and 2) towards the rostral LPFC (bins 4 and 5), task representations become more abstracted from direct perception/action cycles.

### 2.4 Gradient-based meta-analysis of inter-hemispheric asymmetries reveals lateralized associations with topics of language, memory, inhibitory control, and error processing

Our final analysis aims at contrasting the two hemispheres in terms of specific topic associations in a gradient-like fashion. More precisely, we compare homologous quintile bins in both hemispheres in terms of there specific topic associations given unilateral activation. For this purpose, we write a NeuroLang logic program (Program available here) that solves inter-hemispheric segregation queries to infer the probability “*that a topic is present in a study given activation in a right LPFC quintile bin and there exists no reported activation in the entire left LPFC*”. The program also infers the probability of the opposite hypothesis; “*the probability that a Neurosynth topic is present in a study given activation in a left LPFC quintile bin and there exists no reported activation in the entire right LPFC*”. The LOR of these two hypotheses represents the amount of evidence for topic association given unilateral activation in the right hemisphere relative to a unilateral activation in the left hemisphere of the LPFC in a gradient-like fashion.

The results of this analysis are depicted in Figure 5. In general, we do not observe any systematic variation in the degree and nature of hemispheric asymmetries moving along the principal gradient in the LPFC. That is, the amount of evidence for hemispheric dominance as well as the domains of lateralized topic associations in the LPFC are comparable between caudal and rostral LPFC regions, especially in the left hemisphere (see Figure 5). In this context, we find that the left LPFC exhibits greater amount of evidence for topic associations than the right LPFC when given unilateral activations. This result is consistent with findings that the left hemisphere shows a tendency to interact more exclusively with itself than the right hemisphere [50]. Specifically, we find that language-related topics, such as “lexical semantics”, “sentence comprehension”, and “reading” show more than threefold the evidence (*LOR* > 0.5) for left-hemispheric dominance along the entire principal LPFC gradient. Likewise, we find that memory-related topics, “memory retrieval” and “declarative memory”, also show a comparable amount of evidence for left-hemispheric dominance across multiple quintile bins of the principal LPFC gradient as language-related topics. In contrast, topics such as “response inhibition” (in bins 2 and 3), “feedback/error processing” (in bins 2 and 3), and “somatosensory processing” (bins 4 and 5) show weaker evidence (*LOR* < 0.5) for right-hemispheric dominance in the LPFC. Together, these results reassert the views on hemispheric asymmetries in the LPFC, with the left LPFC involved in language and semantic memory processes [22, 30, 33, 51] and the right LPFC involved in stimulus-driven action control and monitoring processes [31, 32, 52, 53].

**Figure 5:**
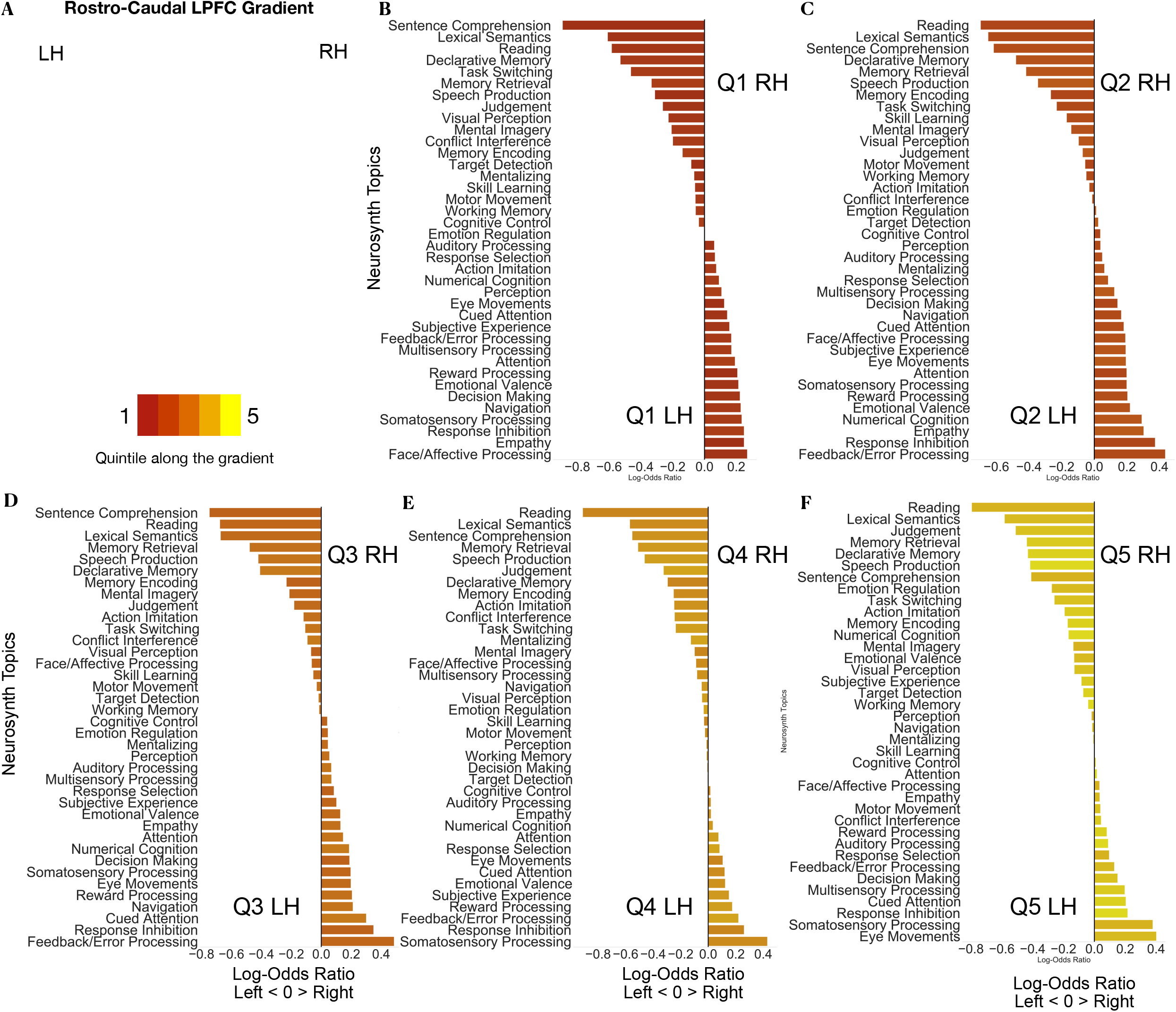
Gradient-based meta-analysis of functional asymmetries reveals the left-hemispheric dominance of language and memory and the right-hemispheric dominance of response inhibition, error processing, and somatosensory processing in the LPFC. Horizontal bar graphs show the topics that are mostly predicted by the presence of unilateral activation in each quintile bin in the right LPFC versus its homologous bin in the left LPFC. Positive log-odds ratios (LORs) indicate evidence in favor of right hemispheric-dominance of a topic in any given bin, whereas negative LORs indicate evidence in favor of left dominance of a topic in any given bin. Topics are ordered from most-left dominant to most-right dominant in each case. **Q**: Quintile, **LH**: Left Hemisphere, **RH**: Right Hemisphere

## 3 Discussion

In this study, we infer the main gradients of organization in the LPFC, a principal rostrocaudal and subsidiary dorsoventral gradient, from a large corpus of literature. These gradients explain most of the variance in meta-analytic connectivity patterns in the LPFC, respectively. We find an agreement in the literature on a spectrum of increasing abstraction both in the spatial distribution of large-scale networks and specific topic associations along the principal gradient in both hemispheres. Finally, when assessing inter-hemispheric asymmetries, we do not observe a systematic transition in the degree of hemispheric-dominance in topic associations along the principal gradient, especially in the left LPFC. Rather we find a pattern of diffusely lateralized topic associations consistent with previous findings on language, memory, and response inhibition. This comprehensive and expressive meta-analysis is enabled by a recently-introduced domain-specific query language, called NeuroLang, that formalizes meta-analysis and expands the scope of hypotheses that can be tested against an ever expanding literature. Overall, the findings of this study can serve to ground new hypothesis generation in future studies on a quantitative overview of previously published results.

### 3.1 The principal gradient of meta-analytic connectivity in the LPFC echoes a rostrocaudal organization characterized by domain-generality in the intermediate zone and domain-specificity at the extremities

The principal gradient of meta-analytic coactivation in the LPFC echoes a rostrocaudal organization in the sense that successive twenty-percentile bins along the gradient show a linear increase in their posterior—to-anterior position from the premotor cortex towards the anterior of the brain. Thus, this gradient places caudal LPFC regions at the farthest point from rostral LPFC regions on a spectrum of similarity in meta-analytic connectivity patterns. This result agrees with a popular class of hypotheses emerging from abstraction and hierarchical control studies on a rostrocaudal gradient in the LPFC [3, 9, 11, 13, 54]. Yet, a question that remains open concerns the properties of different zones along the gradient in terms of domain-generality and domain-specificity.

Early fMRI studies on the organization of the LPFC ascribe the rostral LPFC with the role of integrating concrete information from more caudal regions into more abstract forms and relaying back top-down control signals [2, 11, 13, 54]. However, recent accounts relying on causal evidence argue against a linear gradient, and rather place the mid-LPFC as the nexus of both concrete and abstract representations, with the rostral and caudal LPFC involved in distinct specific domains [6, 8, 29]. Here, we cannot infer such integrative processing by means of causality, nevertheless, we find that the mid-LPFC regions previously described as integrative in [6, 28] overlap with the middle zones (bins 2 and 3) of the principal LPFC gradient. This means that the meta-analytic connectivity profile of mid-LPFC is not completely similar nor dissimilar to those of rostral and caudal LPFC, but rather overlaps with them. Moreover, we find that the coactivation patterns of middle zones (bins 2 and 3) in Figure 3 extends along the LPFC in each hemisphere to include regions of caudal and rostral LPFC, a pattern not observed for the extremities of the gradient (bins 1 and 5). Furthermore, we find that those middle zones predominantly coactivate with the salience (SalVentAttnB) and control networks (ContA and ContB), but to a lesser extent with the attention (DorsAttnA and DorsAttnB) and default mode networks (DefaultA and DefaultB). The salience and control networks are integrative networks believed to mediate the interaction of the default and attention networks to control the transition between internally and externally focused processing according to context [55]. Likewise, this coactivation pattern is not observed for the two extremities of the gradient, where networks involved either in external processing (SomMotA, DorsAttnA, and SalVentAttnA) or internal cognition (DefaultA and DefaultB) are relatively more dominant. Finally, given coactivation restricted within the caudal-to-middle zones of the gradient (bins 1 to 4), we observe associations with topics related to action execution and perception (see Figure 4). In contrast, when coactivation is restricted within middle-to-rostral zones, we observe associations with topics of self-reference, memory, emotion and social cognition. These patterns of network and function associations may indicate domain-generality (i.e. internally and externally oriented processing) in mid-LPFC regions and more domain-specificity (i.e. either internally or externally oriented processing) in caudal and rostral LPFC regions. Thus, the rostrocaudal gradient of meta-analytic connectivity in the LPFC is consistent with the revised view of distinct LPFC hierarchies converging in mid-LPFC [6].

### 3.2 The principal gradient of LPFC meta-analytic connectivity conforms with the principal gradient of brain-wide intrinsic connectivity in both the spatial layout of networks and distribution of functions

The topography of the principal LPFC gradient of meta-analytic connectivity resembles the general layout of the principal brain-wide gradient described in [26], which represents the dominant spatial principle governing the topography of resting-state connectivity throughout the entire cerebral cortex. This spatial principle conceptualizes higher-order cognition as emerging from dynamic interactions of large-scale networks, systematically organized along an axis of abstraction that extends from unimodal sensorimotor regions to transmodal default mode regions [25, 26, 56]. Importantly, it incorporates the seemingly isolated local processing streams across the cortex within a global continuous framework. In this sense, the spatial location of a brain region is not arbitrary; a regions’s position along the principal gradient is a major determinant of its connectivity profile, its network membership and consequently its functional role. Specifically, it has been found that the longer the spatial distance between a region and the primary cortices, the more distant are its functional connections and the more it is dispositioned to subserve abstract mental functions [57]. The default mode network occupies the top end of the principal intrinsic connectivity gradient and exhibits the greatest geodesic distance from the sensorimotor cortices, allowing it to process highly internalized information abstracted from immediate sensory input [26, 58].

In this study, we find that the rostrocaudal gradient in the LPFC captures systematic transitions in large-scale functional networks (Figure 3), such that caudal regions mainly coactivate with sensorimotor/attention networks, middle regions mainly coactivate with salience/executive control networks, and most rostral regions coactivate with the default mode network. Moreover, inferring specific topic associations using NeuroLang’s segregation queries supports the notion of increasing abstraction along the principal LPFC gradient Figure 4. In particular, we find that activations reported only in the caudal end (bins 1 and 2) predict topics of acting and perceiving, while activations restricted to the rostral end (bins 4 and 5) predict topics related to emotion, social cognition, and memory—functions that rely on abstract representations untethered from immediate environmental demands [59]. Interestingly, the presence of the topics “memory retrieval” and “emotion regulation” in both LPFC hemispheres and “response inhibition” in the right LPFC is best predicted by the coactivation between most rostral regions (i.e. quintile bin 5) and more caudal regions (bins 1 to 3, see Figure 4). This result supports a prominent role for the rostral LPFC in retrieving past memories as well as future plans and goals to enable temporal control of behavior and emotions [60]. Ultimately, the rostrocaudal LPFC gradient described herein represents a literature-inferred map of an external/present oriented to internal/temporally-remote organizing spatial principle, wherein globally interacting networks interface locally in the LPFC to support adaptive behavior within dynamic contexts [6, 29].

### 3.3 Segregation-based meta-analysis of inter-hemispheric asymmetries reasserts the left-hemispheric dominance of language and memory and the right-hemispheric dominance of inhibition and feedback/error processing in the LPFC

Segregation-based meta-analysis of inter-hemispheric asymmetries reveals hemisphere-specific associations with language, memory, response inhibition, error-processing, and somatosensory processing in the LPFC. The importance of segregation queries in this case is in inferring the structure-function associations whose presence is predicted by unilateral activation in the LPFC. Previously, the lateralization of function in the brain has been well documented for certain functions, notably language [22, 30] and response inhibition [32]. More recently, an effort to map hemisphere-specific functions across the whole brain [61] has revealed four global dimensions of laterlization: symbolic communication, perception and action, emotion, and decision making. However, a comprehensive comparison of hemisphere-specific functional associations within the LPFC remains lacking, especially when taking into account the principal organizing gradient in each hemisphere. The analysis carried out in this study is one step forward towards filling this gap.

This analysis, however, does not reveal systematic variations in the degree and nature of lateralized structure-function associations along the principal LPFC gradient (Figure 5). That is, unilateral structure-function association patterns seem to be comparable both in topics and strength throughout the gradient, especially in the left hemisphere. The greatest observed evidence for left-hemispheric dominance in the LPFC is attributed to language and memory-related topics, which is consistent with a long line of research on the linguistic and semantic selectivity of the left hemisphere compared to the right hemisphere [34]. In contrast, the greatest amount of evidence for right-hemispheric dominance in the LPFC is attributed to “response inhibition” and “feedback/error processing”, and to a lesser extent “somatosensory processing” and “eye movements”. Surprisingly, we observe relatively weaker evidence (*LOR* < 0.3, see Figure 5) for right-hemispheric dominance of attention-related topics, such as “attention”, “cued attention” and “navigation”, although these are often attributed to the right brain hemisphere [52, 53]. While there may be more than one explanation for these observations, a plausible one is related to the data-driven nature of topics. More specifically, topics are “bags” of words that frequently co-occur in the abstracts of articles, making them at best proxies to the actual mental functions. This means that topics can be noisy and not specific enough to capture finely-grained cognitive constructs. Nonetheless, topics are relatively better representatives of psychological domains than individual terms that pose the risk of being interpreted out of context. Overall, these results support the preferential roles of the left LPFC in language/semantic representations and the right LPFC in sensory monitoring and the cued inhibition of behavior.

### 3.4 Limitations

While the present results provide a relatively unbiased mapping of the organizing gradients in the LPFC through meta-analysis, several limitations are worth noting. First, we make simplifying assumptions in order to improve interpretability and alleviate computational burdens, notably the use of 1024 functional regions from the DiFuMo atlases [43] and the choice of twenty-percentile bins along the gradient as units of analysis. These assumptions might impose a fixed dimensionality on the brain and forces voxels to be grouped in static regions, which ignores dynamics in brain activity observed at multiple timescales within individuals [62]. Another simplifying assumption is the use of topics that represent broad concepts built upon the frequency with which terms co-occur in studies, ignoring more finely-grained cognitive structures. Integrating ontologies, such as the Cognitive Atlas [48], will arguably improve the ability of automated meta-analyses to differentiate fine-grained cognitive constructs. In fact, NeuroLang is well equipped to integrate ontologies into meta-analyses, and this will be our next step towards improving the precision of meta-analytic queries.

A second limitation is that our meta-analysis is based on an automatically-generated coordinate-based dataset. Coordinate-based meta-analytic datasets suffer from information loss due to the relative sparseness of reported results [63], with peak activations being sensitive to statistical methods adopted in each study, notably thresholding [47]. Moreover, with small sample sizes per study, potential “publication bias”, or the tendency of authors and journals to only publish positive or “statistically significant” results [36], might impact the reliability of the current findings. Even though spatial smoothing priors and probabilistic brain atlases may alleviate some bias, future meta-analyses will rely on complete data like unthresholded statistical images stored in large repositories, such as NeuroVault [64], to validate the results.

Finally, an important limitation, not specific to this meta-analysis, is that our current knowledge of task-dependent activation in the brain is as good as the task paradigms that induce these activations [65]. More broadly, an ongoing endeavor in cognitive neuroscience is developing the appropriate paradigms that isolate cognitive processes of closely related brain regions [65]. Studies in the domain of abstraction and hierarchical control use nested tasks classed by different levels of abstraction, which can reveal functional gradients in the LPFC (e.g. [6, 11]). However, these studies are not common in the literature and are limited to small range of functions. In contrast, the bulk of tasks included in the Neurosynth database, while not hierarchical, captures a much wider variety of brain states, but at the expense of losing some level of specificity.

### 3.5 Conclusion

In conclusion, the present study provides quantitative meta-analytic evidence for organizing gradients in the LPFC of humans. The LPFC appears to be organised along two spatial gradients, rostrocaudal and dorsoventral, that respectively explain the most and second-most variance in meta-analytic connectivity. We also reveal that the dominant gradient captures a spectrum of increasing abstraction in network connectivity and specific structure-function associations, grounding a popular class of hypotheses on comprehensive empirical evidence and supporting recent revised views on the functional properties of different LPFC regions. Importantly, we overcome the limitations of previous large-scale attempts using a novel domain-specific query language, called NeuroLang, to formulate expressive queries on the largest coordinate-based meta-analysis database to date. As more studies are aggregated into future databases, the analyses carried out in this study can be reproduced using the same queries as well as extended to explore other brain regions.

## 4 Materials and Methods

### 4.1 Data and Software

We use the latest version of the Neurosynth database [40] last updated in July 2018 to include 14,371 publications with more than 500,000 activation coordinates covering the whole brain. Each study in the database is represented by a PubMed ID, peak activation coordinates and weighted topic associations. Activation coordinates are either reported in MNI space or are transformed from Talairach space before analysis. To examine the structure-function associations in the LPFC, we use the set of 100 Neurosynth topic terms (version 5) previously generated by applying latent Dirichlet allocation to the abstracts of articles in the database [49]. Out of the 100 topics, we include 38 topics that we believe represent coherent cognitive functions, excluding those that correspond to subject populations (e.g. brain disorders, age, sex), brain anatomy, imaging modalities and analysis techniques. Finally, all analyses and visualizations are implemented in python. In particular, we use the NeuroLang (https://github.com/NeuroLang/NeuroLang) package to perform all meta-analysis steps and the BrainSpace package (https://github.com/MICA-MNI/BrainSpace) to estimate a low-dimensional embedding of meta-analytic connectivity patterns in the LPFC [46]. All python notebooks and data files used in this study will be publicly available to be openly accessed and used on https://osf.io/ur7ej/quickfiles.

### 4.2 The lateral prefrontal cortex mask

To facilitate the selection of regions in the LPFC for meta-analysis, a spatial mask of the LPFC is needed. We rely on a previously created mask of the lateral frontal lobe created from [9]. However, we exclude voxels with less than 25% probability of falling in the grey matter as well as voxels located at *x* < 18 or *x* > −18 from the midline of the brain to ensure that regions in the anterior and superior parts of the medial prefrontal cortex are not included. We also exclude voxels in the orbitofrontal cortex and anterior insula, while making sure to include voxels of the lateral orbitofrontal cortex. Finally, to focus our analysis on the association regions of the lateral frontal lobe (i.e, the LPFC), we exclude voxels in the motor cortex as defined by the somatomotor networks of the 17-Networks atlas [24]. The LPFC mask is shown in Figure 6A.

**Figure 6:**
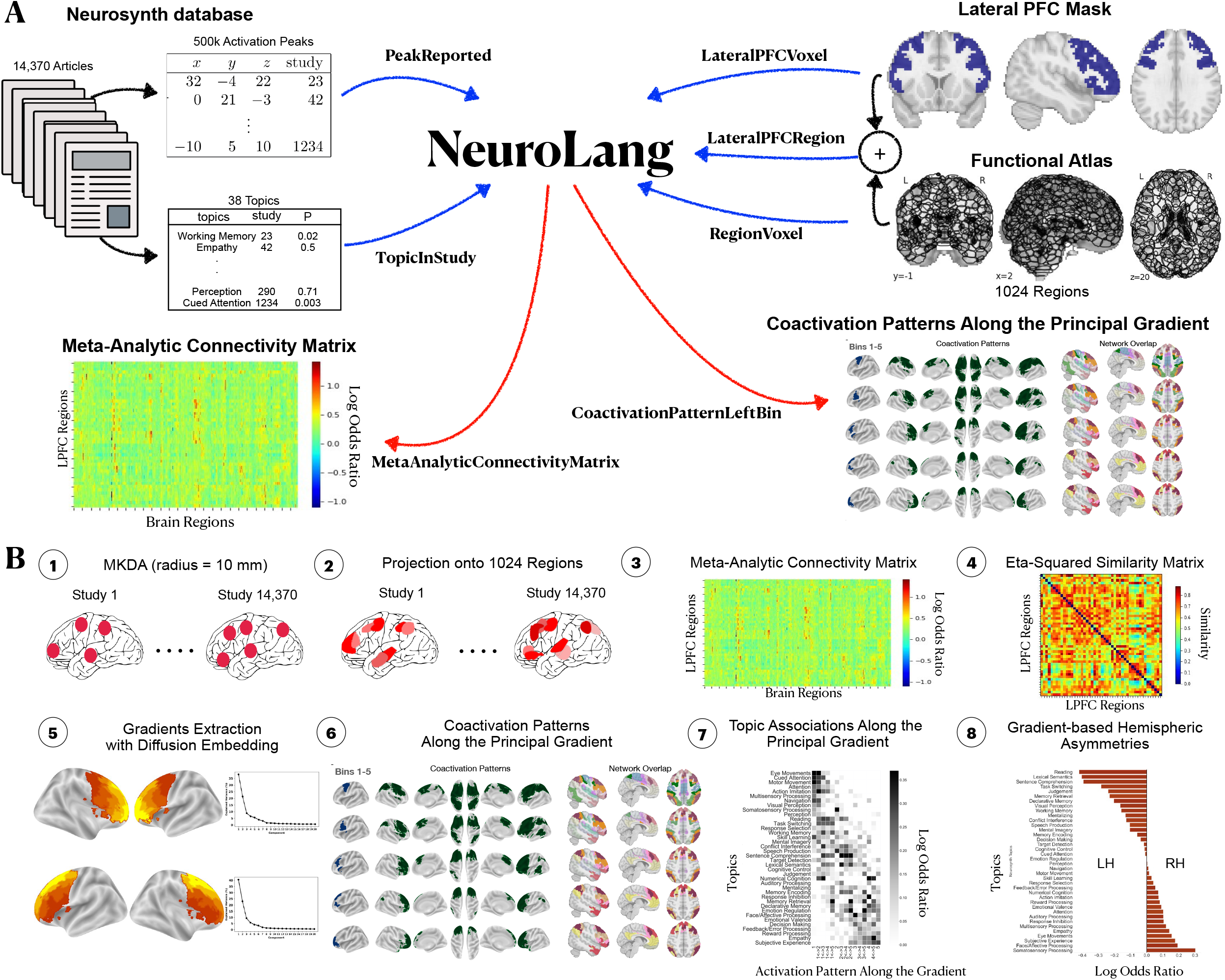
Schematic overview of our analysis pipeline. **(A)** Inputs and outputs of NeuroLang. Inputs are represented using blue arrows and include: Peak activations and topics from the Neurosynth database, the lateral PFC mask, and the 1024 regions from the DiFuMo atlases are represented in a unifying framework within NeuroLang. Two examples of outputs are shown here and represented using red arrows. **(B)** The main steps of the meta-analysis carried out in this study. (1) Multilevel Kernel Density Analysis (MKDA) selects voxels within 10 mm radius of each peak activation in each study for meta-analysis. (2) The binary activation map of each study is projected onto 1024 functional regions. Varying shades of red signify that regions have different probabilities of being reported by a study depending on the location of reported voxels with respect to each region. (3) The meta-analytic connectivity matrix is inferred, and it encodes the log-odds ratios of coactivation between each region in the LPFC and every region in the brain. (4) The degree of similarity between LPFC regions in their meta-analytic connectivity profiles is estimated via eta-squared similarity metric. (5) The gradients of meta-analytic connectivity in each hemisphere are derived from the affinity matrix using Diffusion Embedding. (6) Coactivation patterns of successive quintile bins along the gradient are inferred, as well as their overlap with large-scale networks. (7) Specific topic associations along the principal gradient are inferred using segregation queries. (8) Finally, gradient-based metaanalysis of hemispheric asymmetries is performed using segregation queries between homologous quintile bins.

### 4.3 The 1024 functional regions dictionary from DiFuMo

To increase the interpretability of our findings and alleviate computational burdens, we reduce voxel-level data to region-level data. In particular, we adopt the 1024 functional regions dictionary from the Dictionaries of Functional Modes (DiFuMo) atlases [43]. The DiFuMo is a set of multi-scale functional atlases estimated via massive online dictionary learning [66] applied to functional brain volumes of thousands of subjects across 27 large-scale studies, forming a total of 2192 task-based and resting-state MRI sessions. Reducing voxel data to 1024 functional regions has been argued to capture the functional neuroanatomy of the brain equally well as voxel-level analysis while reducing computational burdens Dadi et al. [43]. Unlike other dimensionality-reduction techniques, massive online dictionary learning assigns non-negative continuous loadings to each voxel designating its relative weight on each region. Voxels that have a loading value equal to 0 on any given region are considered to not belong in this region. Finally, to identify the regions in the LPFC, we recover those that have at least 50% of their volume fall within the LPFC mask described earlier (see the next section on Representing heterogeneous data in a single framework with NeuroLang). Note that we do not mask out voxels of regions that are outside the LPFC mask—we either include or exclude entire regions without breaking continuity. This means that some functional regions can include voxels outside the LPFC mask. The reason for this crossover is that functionally-defined regions seldom conform to anatomical landmarks in the brain. Comprehensive details on DiFuMo can be found in the original study by Dadi et al. [43].

### 4.4 Representing heterogeneous data in a single framework with NeuroLang

The goal behind developing NeuroLang is to create a universal language that reduces the likelihood of miscommunication within the cognitive neuroscience community by enabling databases, hypotheses, and questions to be defined in a formal, shareable and reproducible manner. This is believed to be a critical step towards advancing the field of cognitive neuroscience [65].

In this study, we represent various data types coming from heterogeneous sources, such as peak coordinates, topic models, anatomical masks, and brain atlases in a single framework. More precisely, these data and the relationships among them can be represented as facts and rules using declarative logic-based statements such that the user only has to specify what is to be found rather than how to find it. Facts and rules in NeuroLang are tuple-sets or tables structured in rows. Each row is a sequence of *k* elements representing a piece of data, such as the MNI coordinates of a reported peak in a study, and can be implicitly assigned a probability that quantifies the level of uncertainty in this data. Fact tables represent explicit information present in the data, while rule tables represent inferred relationships among the different data elements. The goal is to declare these tables as **predicates** in a probabilistic logic program that solves complex queries on them. For a survey on probabilistic databases and probabilistic programming the reader is referred to [67].

To concretely showcase how we represent data in NeuroLang, we start with the Neurosynth database. the database includes studies that report peak activations coordinates in standard space (Figure 6A). In NeuroLang, we represent these peaks in a fact table called `PeakReported`. This table contains a row `(x, y, z, study)` for each peak with coordinates `(x, y, z)` that has been reported active by a `study`. Also, the studies themselves are represented in a fact table called `Study` that contains a row for each study containing a single element, `(study)`, representing its PubMed identifier. Similarly to Neurosynth, we assume each study within the database to be an *independent equiprobable sample* of neuroscientific knowledge [40]. This assumption is represented by another fact table we call `SelectedStudy`, which simply assigns a uniform probability (1/*N*, *N* = 14, 370) for each study to be selected in any possible world of events. In other words, this assumption allows the studies to have a similar weight in the meta-analysis [40, 42].

Further, the spatial uncertainty surrounding the reported location of each peak in a given study can be represented in a rule table, named `VoxelReported`. In this rule, a multilevel kernel density analysis (MKDA) [47] assumes each peak’s 10mm neighboring voxels to be equivalently reported [47]. Then, the `VoxelReported` table contains a row `(x, y, z, study)` for each voxel at location `(x, y, z)` and falls within 10 mm Euclidean distance from a peak reported by a `study`. Being based on Datalog [68], a fully declarative logic programming language designed to solve queries on large databases, the NeuroLang program that computes `VoxelReported` is written as follows:

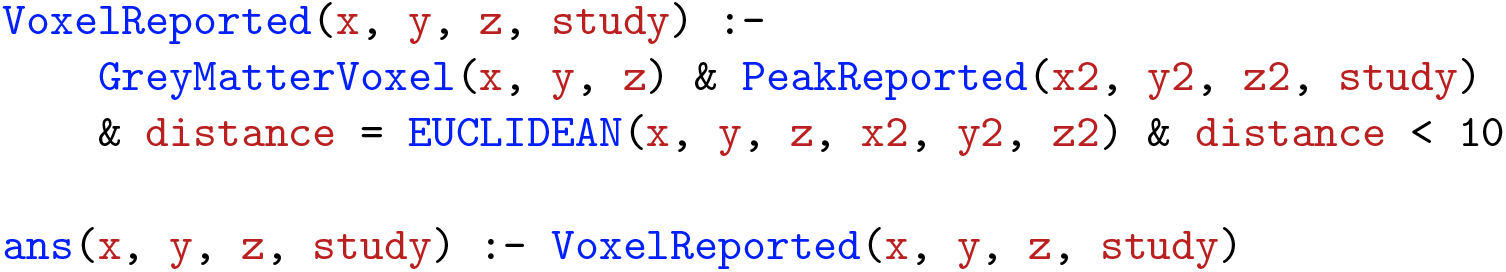

The answer states that “a voxel at location `(x, y, z)` in grey matter is considered active in a `study` if it is situated within 10 mm radius from a peak reported by `study` at location `(x2, y2, z2)`”. Here, `GreyMatterVoxel` is also a fact table representing a grey matter mask in MNI space. This table contains a row `(x, y, z)` for each brain voxel having at least 25% chance of being found in grey matter. The `distance` variable is estimated using the built-in function `EUCLIDEAN`, which computes the distance between two locations in standard space.

The Neurosynth database also includes topics that have been derived using latent Dirichlet allocation applied to the abstracts of the articles [65]. Each study in the database has a loading value on each topic, which can be considered as a proxy to the probability that a topic is present in a study. This weighted topic-study association is represented in a probabilistic fact table named `TopicInStudy`. This table contains a row `(topic, study)` for each `topic` present in a `study`, and the study has a non-zero loading on the topic. The reason for calling this table probabilistic is that the study-on-topic loading is implicitly embedded as a measure of uncertainty in the presence of a topic in a given study.

Anatomical masks and functional atlases can also be represented in NeuroLang. For instance, we represent the LPFC mask described previously in a fact table called `LateralPFCVoxel`. This table contains a row `(x, y, z)` for each voxel belonging to the LPFC mask. Moreover, we represent the 1024 functional regions from DiFuMo in a fact table called `RegionVoxelWeighted`, which contains a row `(r, x, y, z, w)` for each voxel at location `(x, y, z)` in MNI space having a non-zero weight `w` on a DiFuMo region `r`. Similar to case of topic-study association, the voxel-on-region weight can be used as a measure of uncertainty in a voxel belonging to a region. This is achieved by first scaling the weight of every voxel in each region to the maximum weight value in that region. In this sense, the voxel with the maximum loading on a region will have a probability of 1 of belonging to it. This creates a probabilistic fact table `RegionVoxel`, named this way as it does not contain the weight variable explicitly, but implicitly. This probabilistic table can be used by a NeuroLang program to infer other probabilistic tables. This will be concretely shown in the following sections. Finally, regions belonging to the LPFC can be represented in a rule table called `LateralPFCRegion`. This table contains a row `(r)`, where `r` is a brain region that have at least 50% of its volume overlapping with the LPFC mask. The following NeuroLang program, written in Datalog syntax, infers this table:

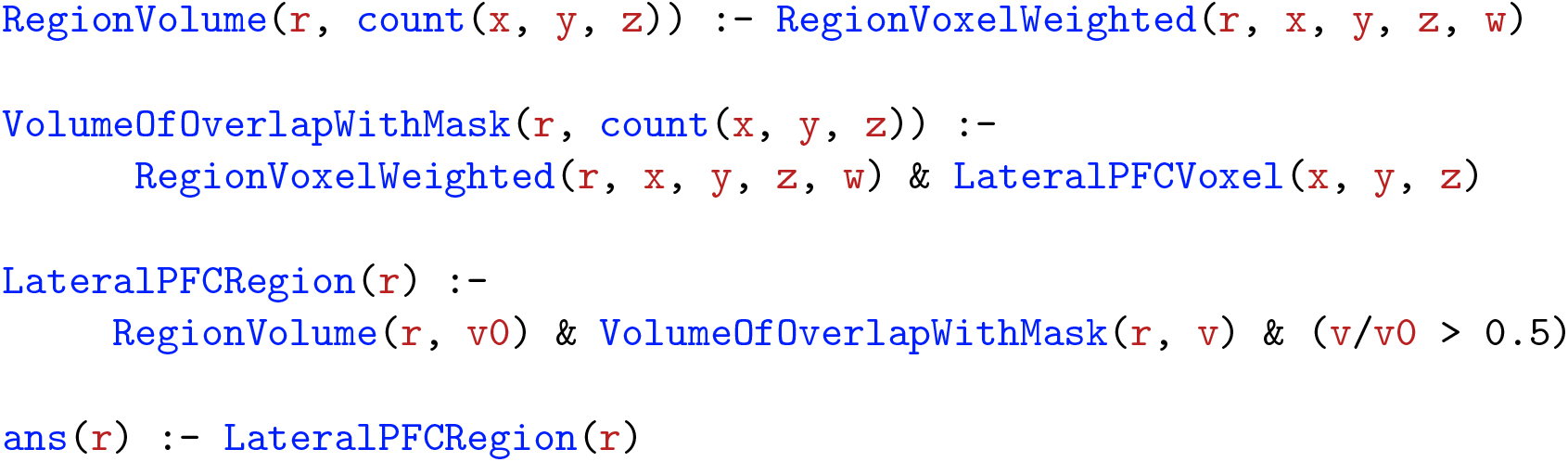

We start by declaring the predicates to be used in solving the query. These predicates are the `RegionVolume` and `VolumeOfOverlapWithMask`. These encode the total volume and the volume of overlap with the LPFC mask of each brain region `r`, respectively. The `LateralPFCRegion` is the final answer of this program and states that: “A brain region `r` belongs to the LPFC if its volume of overlap, `v`, with an LPFC mask makes up more than 50% of its total volume, `v0`”. Volume variables `v` and `v0` are estimated by the built-in function `count(x, y, z)`, which simply counts the number of voxels in a brain region.

### 4.5 Inferring the meta-analytic connectivity matrix using NeuroLang

To infer a whole-brain meta-analytic connectivity profile for each LPFC region, we query the database on the probability that a brain region is reported active given the presence as well as when given the absence of activation in a LPFC region. This gives us a measure of specificity in the meta-analytic connectivity between each LPFC region and every brain region.

To infer these probabilities, we write a NeuroLang program that first projects the voxels reported active in each study onto the 1024 functional regions to determine which ones are reported by the study (step 2 in Figure 6B). In this context, the program regards the reporting of a brain region by a study as a probabilistic event rather than a certain one. That is, if a voxel reported active has a normalized weight `w` on a region `r`, then the region is assigned a probability `w` of being reported by the study. If multiple voxels are reported active within a region, then the union of their locations’ weights is considered the overall probability for the region to be reported by the study. This union is interpreted as the probability that at least one of those peaks is reported active in the region. Intuitively, a region is considered to be not reported by a study, if no activation is reported in any of its constituent voxels.

We then infer the conditional probabilities of observing activation in a brain region given the presence, and subsequently given the absence, of activation in a LPFC region. To quantitatively contrast these two hypotheses and get a representative measure of meta-analytic connectivity, we compute the logarithm with base 10 of their odds ratio (LOR). This yields a vector for each LPFC region whose elements represent the amount of evidence (in log-scale) for pairwise meta-analytic connectivity with every brain region. A positive LOR indicates more evidence for a brain region to be reported active given activation in a LPFC region, a negative LOR implies more evidence for a brain region to be reported active given no activation in the LPFC region, and a LOR equal to 0 implies that the evidence is inconclusive for either hypotheses. The program that infers the meta-analytic connectivity matrix is as follows:

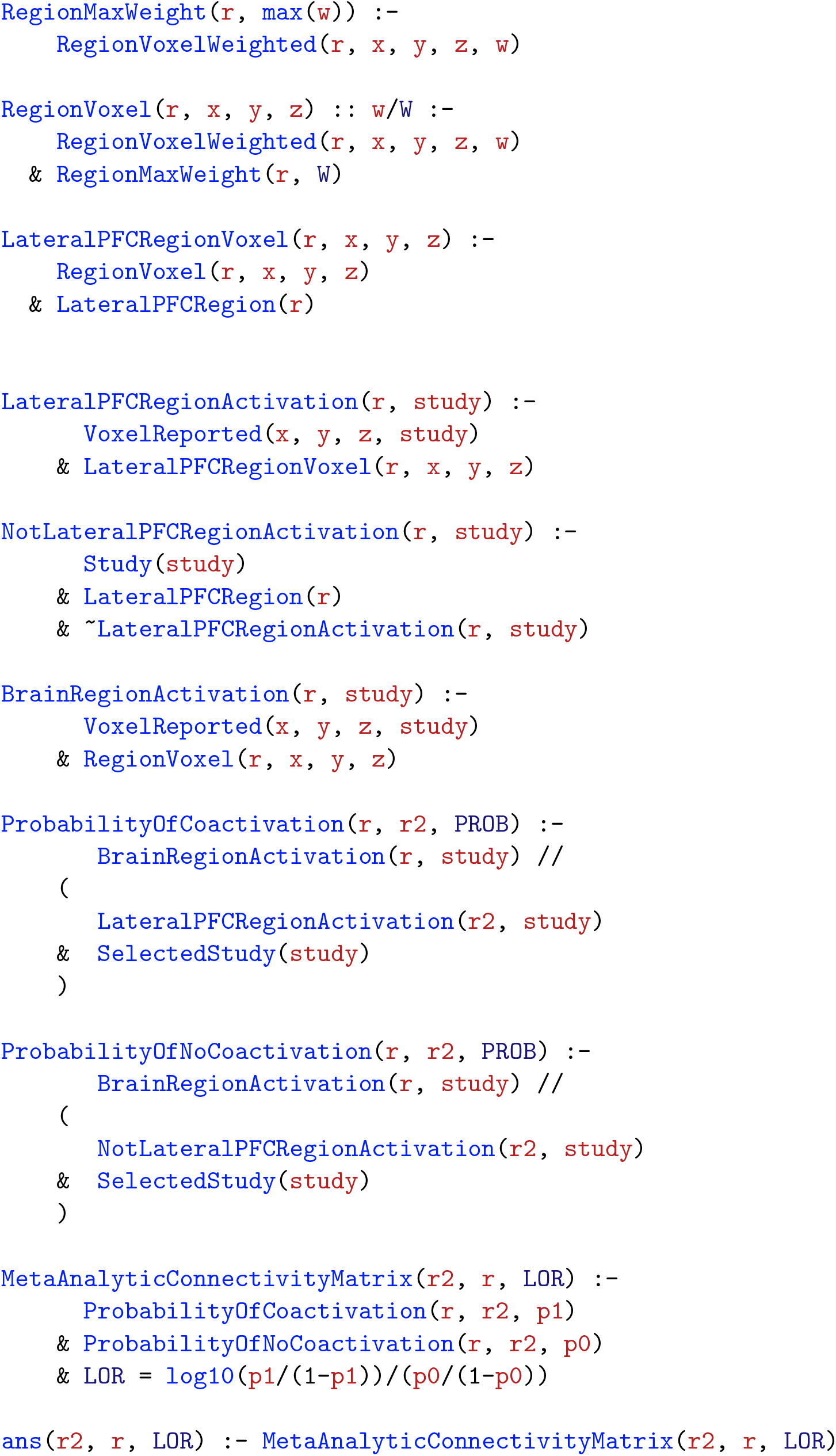

In order for the program to solve the query, we need to declare some useful predicates. First, we get the maximum weight in each DiFuMo brain region in `RegionMaxWeight` using the built-in function `max(w)`. This will be used to declare the probabilistic table `RegionVoxel` which implicitly incorporates the normalized weight `w/W` of each voxel `(x, y, z)` on each region `r`. This is represented by the `(:: w/W)` notation after `RegionVoxel(r, x, y, z)`. Second, we define the tables `LateralPFCRegionActivation` and `BrainRegionActivation`. These are probabilistic rule tables, wherein each row is implicitly assigned a probability that a given brain or LPFC region is reported active by a `study`. Likewise, we declare the studies that do not report activation in each LPFC region by using the negation sign **“**~**”** before `LateralPFCRegionActivation`. Each row of this table represents a `study` that does not report activation in a LPFC region with a certain level of uncertainty. To be able to obtain this table in safe range, we must re-assert that the variable `study` comes from the fact table `Study` and the variable `r` is found in the fact table `LateralPFCRegion`.

To infer the conditional probabilities, we use the “`//`” sign, which means “given”. For instance, `ProbabilityOfCoactivation` is a rule table that encodes the probability (`PROB`) of activation being reported in brain region `r` **given** that activation is also reported in LPFC region `r2`. Likewise, `ProbabilityOfNoCoactivation` is another rule table that encodes the probability (`PROB`) of activation being reported in brain region `r` **given** that activation is **not** reported in LPFC region `r2`. The `SelectedStudy` table sets the program to assign an equal weight (1/*N*, *N* = 14, 370) to all the studies in the meta-analysis. Finally, the `MetaAnalyticConnectivityMatrix` rule table is inferred by computing the “`LOR`” of the two hypotheses as a measure of evidence of meta-analytic connectivity between each LPFC region and every brain region.

### 4.6 Diffusion map embedding using the BrainSpace toolbox

To recover a low-dimensional embedding of the meta-analytic connectivity matrix, we choose to apply diffusion embedding [45], an unsupervised nonlinear dimensionality reduction method. The low-dimensional embedding reveals the axes of variation in coactivation-based connectivity patterns in the LPFC, and can be recovered with two steps. First, we estimate the similarity between LPFC regions in terms of their coactivation patterns. Here, we quantify similarity between each pair of LPFC regions using the eta-squared coefficient following [69], yielding a square affinity matrix (step 4 in Figure 6B). The eta-squared coefficient represents the fraction of the variance in one meta-analytic connectivity profile that is accounted for by the variance in another, and ranges from 0 (totally dissimilar) to 1 (perfectly similar). Diffusion embedding then represents this similarity structure as an arrangement of regions in an embedding space spanned by 20 components known as “gradients”. Gradients are conceptually similar to the components of principal components analysis and represent unidimensional axes each explaining a fraction of the variance in a given feature [26], in our case, meta-analytic connectivity. In each gradient, regions that have very similar meta-analytic connectivity patterns occupy nearby zones, while regions with dissimilar patterns are situated further apart. The first or principal gradient is the most informative component as it captures the dominant axis of variation in meta-analytic connectivity patterns within the LPFC.

### 4.7 Inferring whole-brain coactivation patterns of quintile bins along the principal gradient using NeuroLang

To be able to infer varying coactivation patterns along the principal gradient in the LPFC, we first create regions-of-interest from successive twenty-percentile bins (i.e. five quintile bins) along the gradient in each hemisphere. Then, we identify the large-scale brain networks that overlap with each quintile bin’s coactivation pattern to characterize the variation of network connectivity along the principal gradient (step 6 in Figure 6B). We represent the regions-of-interest created from quintile bins in each hemisphere as fact tables called `LeftBinVoxel` and `RightBinVoxel` for the left and right LPFC, respectively. Each of these tables includes a row `(bin, x, y, z)` for each voxel at location `(x, y, z)` in MNI space and belonging to a quintile `bin`. Moreover, we declare another fact table `Bin` whose rows contain only the labels of the quintile bins (i.e., bin1 to bin5).

We write a NeuroLang program that infers the conditional probability of a brain region to be reported active given activation reported in a bin as well as when given no bin activation. Specifically,the program infers the LOR of these two hypotheses as a measure of evidence for coactivation between each brain region and each quintile bin along the principal gradient. A cortical coactivation pattern for each quintile bin is then constructed by recovering the brain regions that exhibit at least threefold the evidence (or `LOR > 0.5`) of being reported active when given activation in a bin relative to no bin activation. The NeuroLang program that infers coactivation patterns of quintile bins in the left LPFC is as follows:

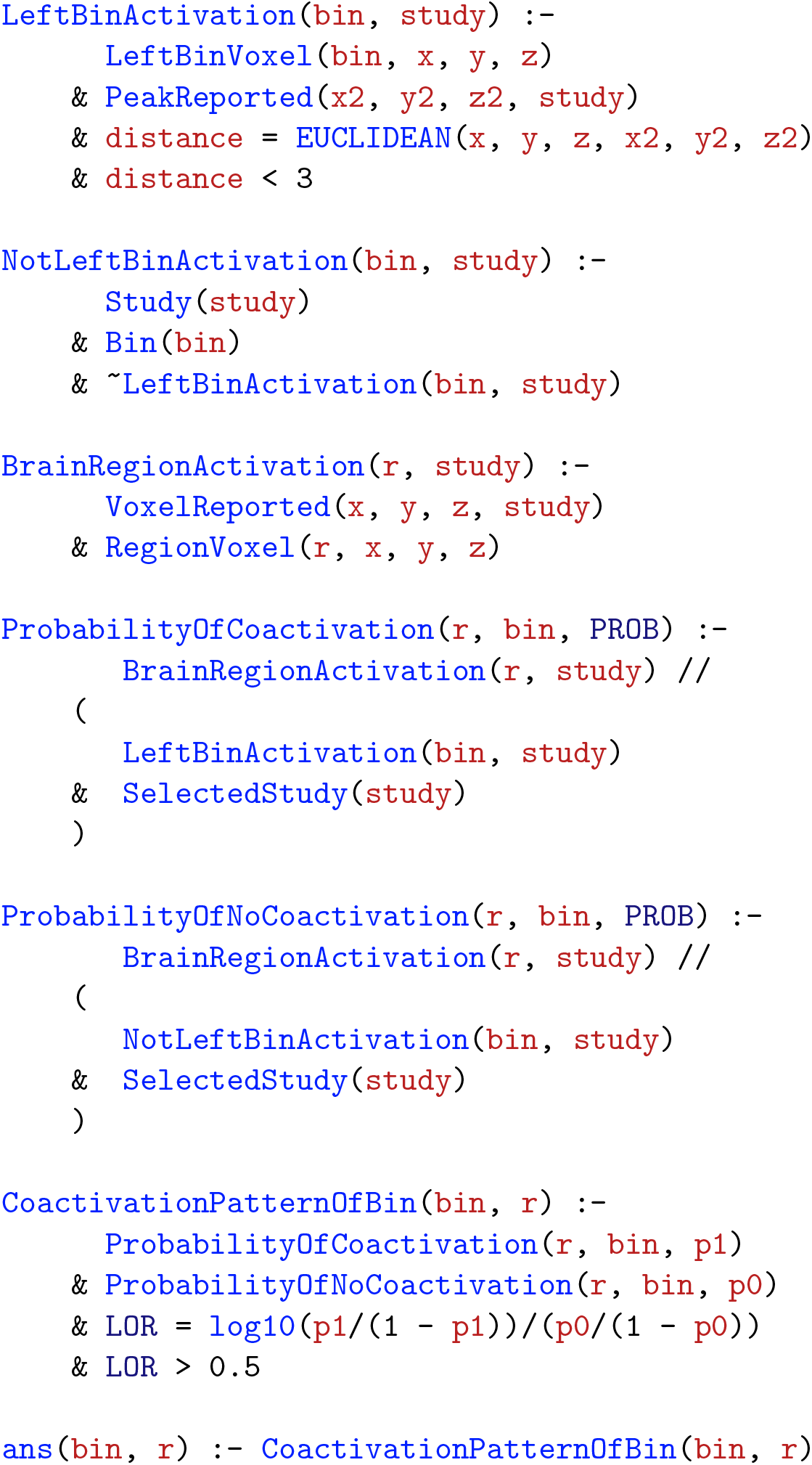

As in the program of the previous section, we first declare the predicates that will be used to find the answer to our query. We set the program to consider activity in a quintile bin to be reported by a study if at least one peak activation is reported within the bin or within its near vicinity (`distance < 3`). The program stores the results in the rule table `LeftBinActivation`, which includes a row `(bin, study)` for each `bin` in which activation is reported by a `study`. The program then derives the studies that do not report activity within each bin using the negation operator and stores them in another rule table `NotLeftBinActivation`. This rule table includes a row for each `bin` wherein no activation has been reported by a `study`. Similarly as the program in the previous section, the program here considers activation reporting in individual brain regions as a probabilistic rather than deterministic event depending on the location of active voxels within each region, and stores the results in the rule table `BrainRegionActivation`. The program then infers the conditional probabilities of the two hypotheses and stores them in the rule tables `ProbabilityOfCoactivation` and `ProbabilityOfNoCoactivation`. Finally, the answer to our query `CoactivationPatternOfBin` is derived by estimating the “`LOR`” as a measure of evidence in favor of coactivation between each brain region `r` and each `bin` and thresholding it at `LOR > 0.5`. Below is a similar program that infers coactivation patterns of quintile bins in the right LPFC:

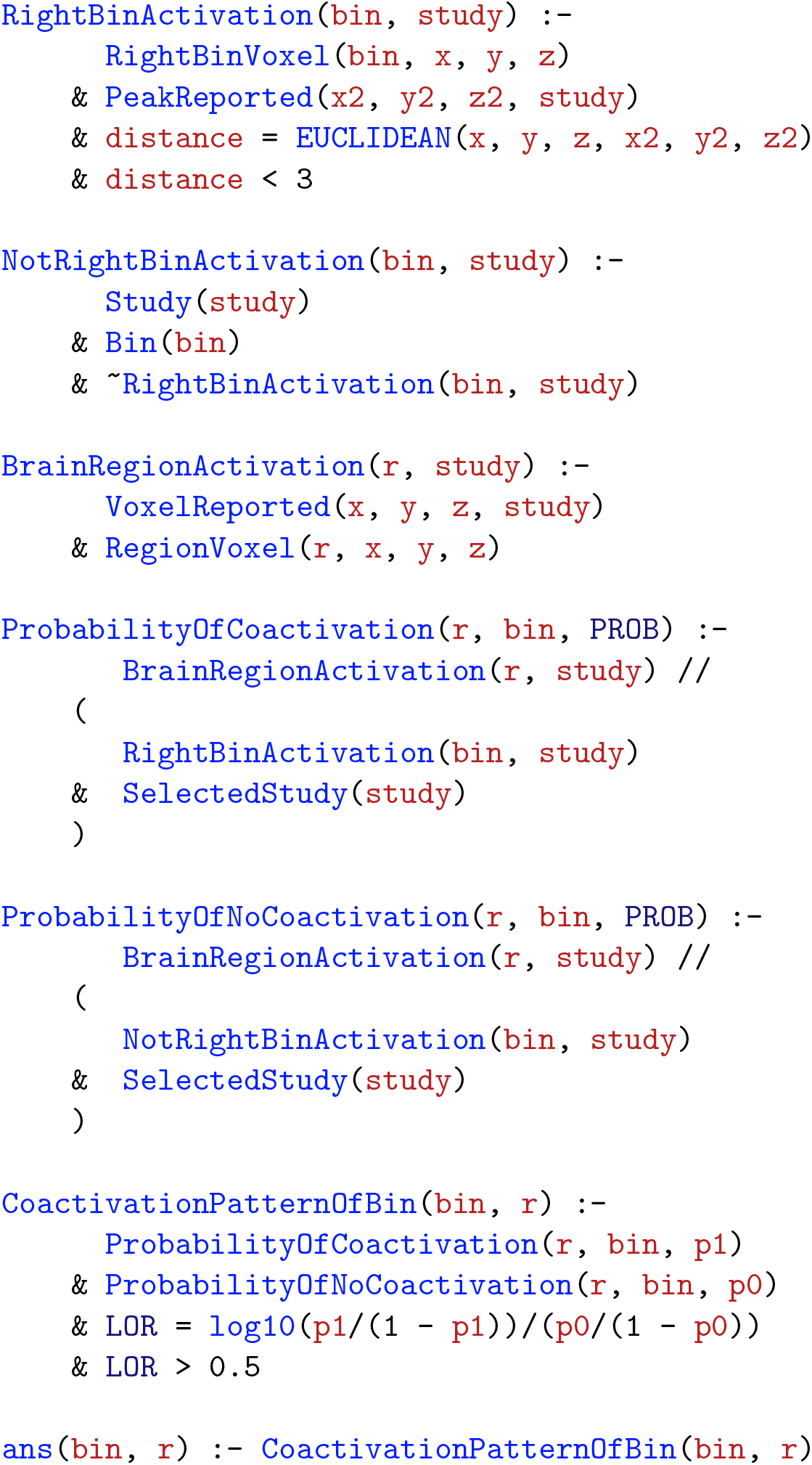

### 4.8 Inferring specific structure-function associations using NeuroLang segregation queries

We infer specific structure-function associations by estimating the extent to which a spatially-localized activation along the principal gradient in the LPFC predicts a Neurosynth topic’s presence in a study. For this purpose, we write NeuroLang programs (see below) that includes what we call “segregation queries”.

Intra-hemispheric segregation queries infer “the probability that a topic is present in a study given spatially-constrained activation within a range of quintile bins and the **simultaneous absence of activation outside this range within the same hemisphere**”. Concurrently, a segregation query infers the probability of the opposite hypothesis: a topic is present given no activation within the range of quintile bins **or** there exists activation outside the range. The LOR of these two hypotheses gives us a measure of evidence in favor of association between a topic and patterns of activity along the principal gradient. The NeuroLang program that infers specific structure-function associations in the left LPFC using segregation queries is as follows:

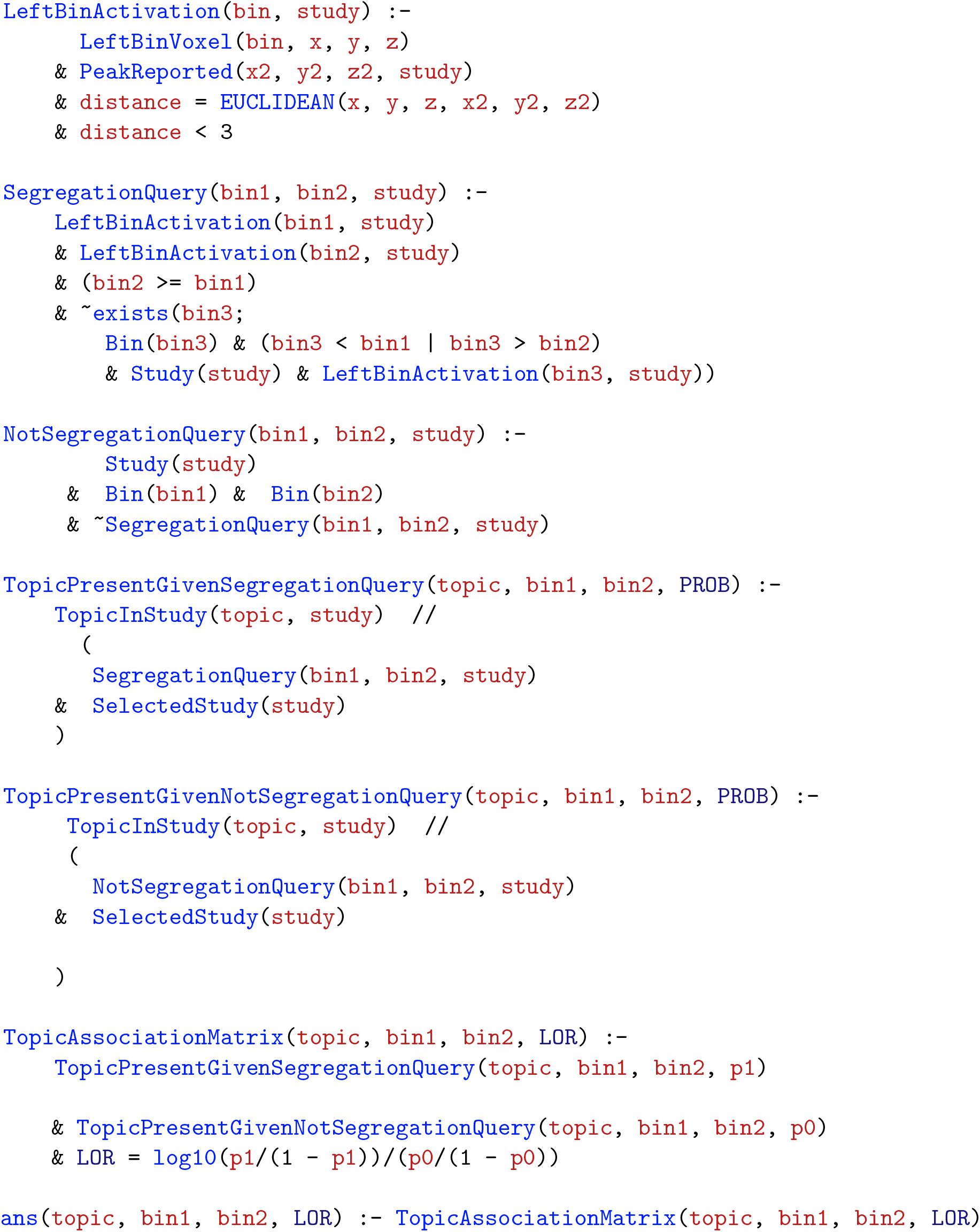

We first declare the studies that report activations in each quintile bin of the left LPFC principal gradient in a rule table `LeftBinActivation`. Then, we declare a segregation query which first identifies the studies that report coactivation between each pair of quintile bins along the principal gradient, `bin1` and `bin2`, under the conditions that `(bin2 >= bin1)` **and there exists no** activation reported in any `bin3`, such that `(bin3 < bin1 | bin3 > bin2)`. That is, activation in any bin that is outside the range `[bin1, bin2]` should not be present, whereas activation in the range between the bins can exist. Here, “**there exists no**” is represented by ^~^exists, which is a combination of the negation operator and the existential quantifier. The results are represented in a `SegregationQuery` rule table, which includes a row `(bin1, bin2, study)` for bins `(bin1, bin2)` between which activation is reported in `study` that also satisfies the segregation condition. Concurrently, we declare the studies that do not match the conditions of the segregation query, and represent them in the rule table `NotSegregationQuery`.

After defining the useful predicates, the program infers the conditional probability that a `topic` is present in a `study` given the presence as well as absence of the segregation condition. The results are represented in the tables `TopicPresentGivenSegregation` and `TopicPresentGivenNotSegregation` Finally, the answer to our query, represented in the rule table `TopicBinsAssociationMatrix`, is derived by computing the “`LOR`” of the two hypotheses as a measure of evidence in favor of specific association between each topic `topic` and activation in a quintile range `[bin1, bin2]` along the principal gradient. To ensure that the results are not driven by a single choice of studies, we run this NeuroLang program 1000 times using random sub-samples of the Neurosynth database (80% of the dataset) in each run. This procedure creates an empirical distribution for each probability estimation from which we consider the 95^*th*^ percentile as a point estimate of interest. The NeuroLang program that infers specific topic associations of coactivation patterns within the right LPFC is as follows:

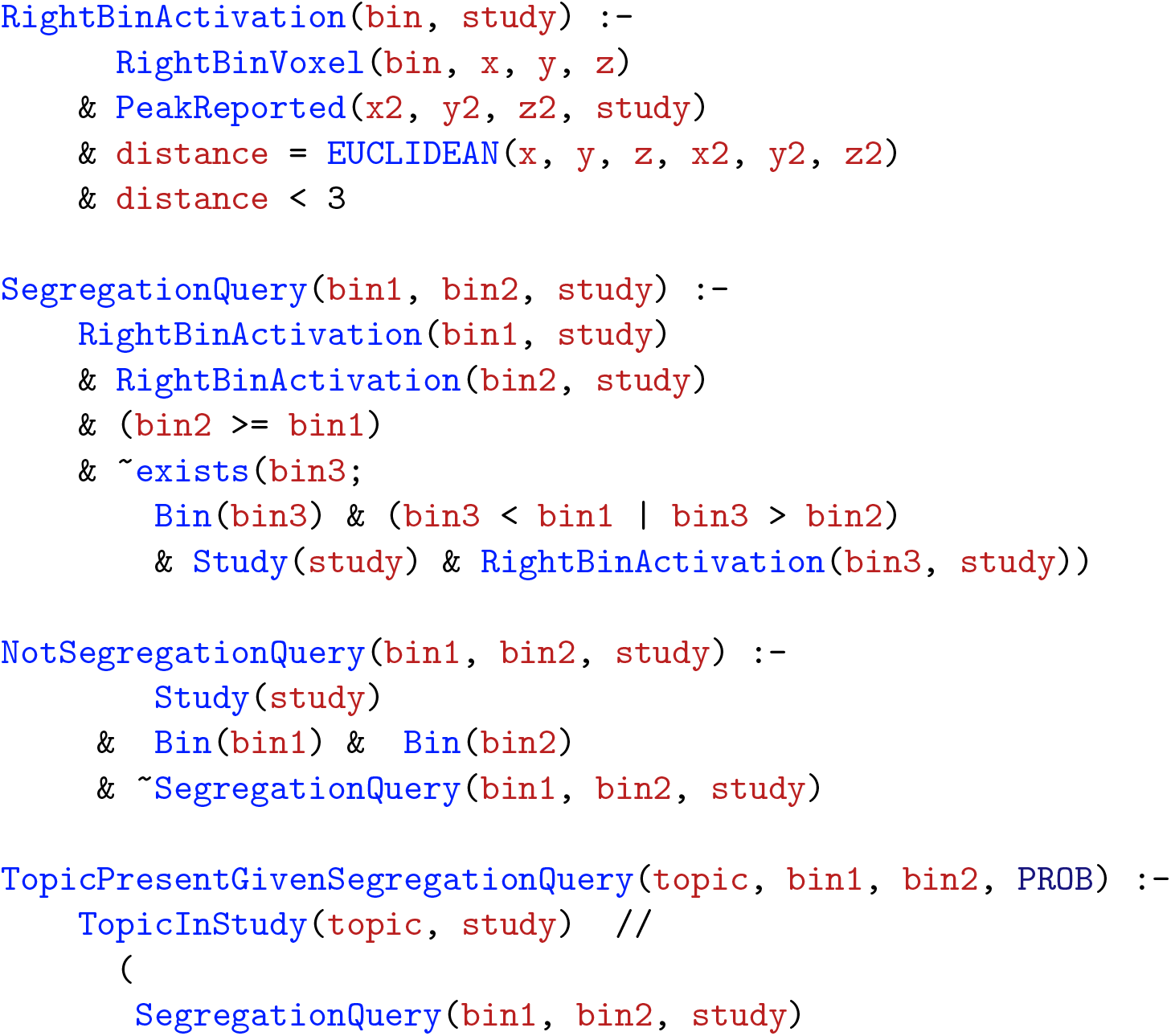
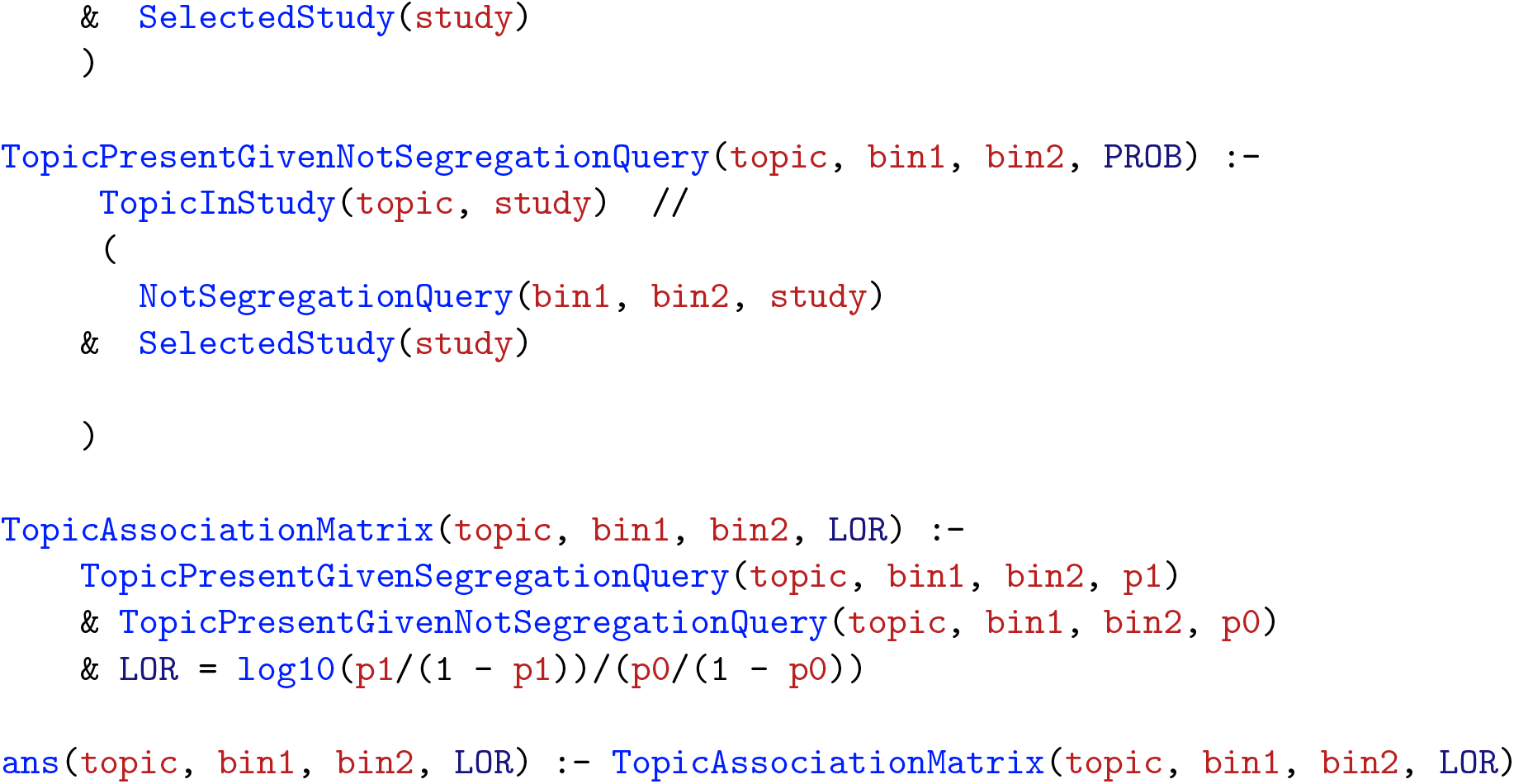

Finally, we write a program that performs inter-hemispheric segregation queries to infer the probability “*that a topic is present given activation in a right LPFC quintile bin and there exists no reported activation in the entire left LPFC*”. The program also infers the probability of the opposite hypothesis; “*a topic is present given activation in a left LPFC quintile bin and there exists no reported activation in the entire right LPFC*”. The NeuroLang program that infers hemisphere-specific topic-bin associations is as follows:

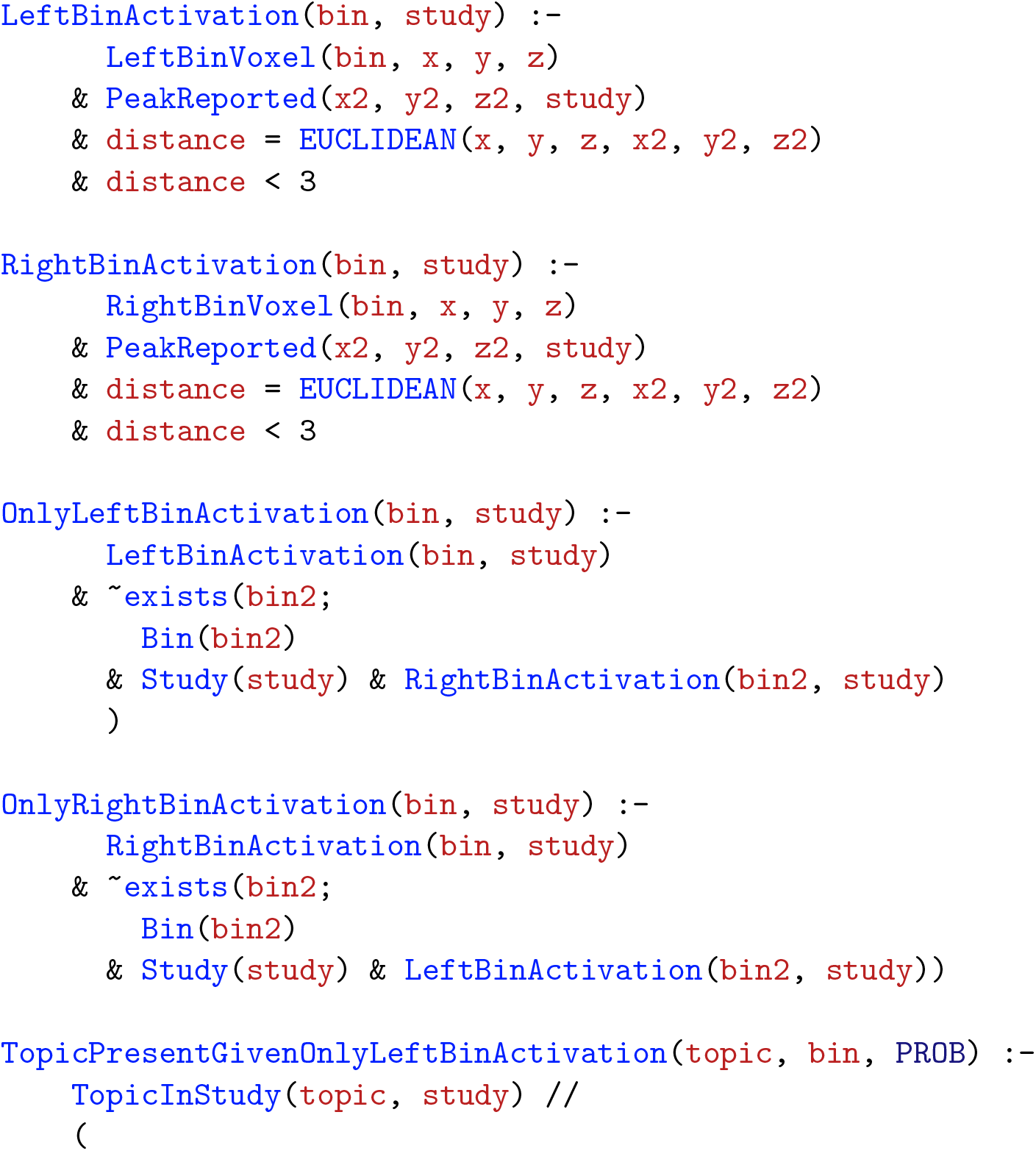
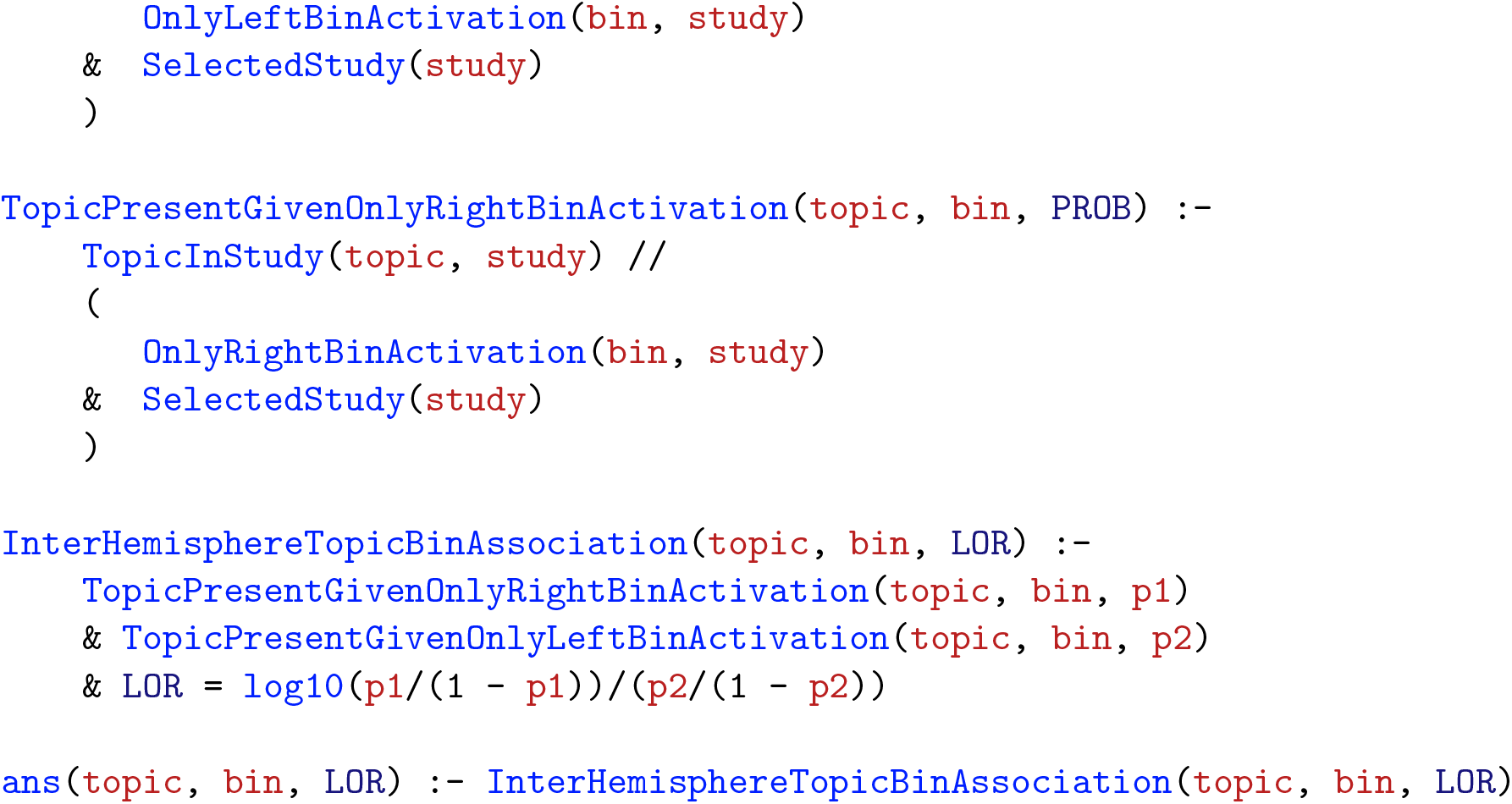

In this program, we define the predicates `LeftBinActivation` and `RightBinActivation` that store the studies reporting activation in each quintile bin of the principal gradient in the left and right LPFC, respectively. Then we declare the inter-hemispheric segregation queries using the negation operator and the existential quantifier, ^~^exists, and stores the results in `OnlyLeftBinActivation` and `OnlyRightBinActivation`. Subsequently, the program infers the conditional probabilities that a `topic` is present in a `study` when given activation either in a left or a right quintile bin. The final answer, `InterHemisphereTopicBinAssociation` is derived by computing the “`LOR`” of the two hypotheses. We run this NeuroLang program 1000 times using random sub-samples of the Neurosynth database (80% of the dataset) in each run, which yields an empirical distribution for each conditional probability estimation from which we consider the 95^*th*^ percentile as the point estimate of interest.

## 5 Data Availability Statement

All data and scripts used in this study are openly available to be accessed and freely used by the community. The source code of NeuroLang is freely available on GitHub at https://github.com/NeuroLang/NeuroLang.

## 6 Acknowledgements

This work is funded by the ERC-2017-STG NeuroLang grant. We are grateful to Jonas Renault who worked on optimising NeuroLang’s engine and develop its web interface.

